# An Improved Human smORF Annnotation Workflow Combining De Novo Transcriptome Assembly and Ribo-Seq

**DOI:** 10.1101/523860

**Authors:** Thomas F. Martinez, Qian Chu, Cynthia Donaldson, Dan Tan, Maxim N. Shokhirev, Alan Saghatelian

## Abstract

Protein-coding small open reading frames (smORFs) are emerging as an important class of genes, however, the coding capacity of smORFs in the human genome is unclear. By integrating *de novo* transcriptome assembly and Ribo-Seq, we confidently annotate thousands of novel translated smORFs in three human cell lines. We find that smORF translation prediction is noisier than for annotated coding sequences, underscoring the importance of analyzing multiple experiments and footprinting conditions. These smORFs are located within non-coding and antisense transcripts, the UTRs of mRNAs, and unannotated transcripts. Analysis of RNA levels and translation efficiency during cellular stress identifies regulated smORFs, providing an approach to select smORFs for further investigation. Sequence conservation and signatures of positive selection indicate that encoded microproteins are likely functional. Additionally, proteomics data from enriched human leukocyte antigen complexes validates the translation of hundreds of smORFs and positions them as a source of novel antigens. Thus, smORFs represent a significant number of important, yet unexplored human genes.

---

The early annotation of open reading frames (ORFs) from nucleic acid sequencing was carried out by locating in-frame start (AUG) and stop codons^1–4^. This approach resulted in unreasonably large numbers of ORFs < 100 codons called small open reading frames (smORFs)^1,3,4^. A length cutoff was then introduced to remove smORFs^5,6^, which were largely presumed to be meaningless random occurrences^1,3^. With the advent of more sensitive detection methods, functional proteins encoded by smORFs, dubbed microproteins, have been characterized with more regularity^7–9^. Early on, gene deletion experiments in yeast identified a number of smORFs that control growth under different environmental conditions^10^. In fruit flies, a gene called tal/pri was identified, which contains four smORFs encoding three 11- and one 32-amino acid microproteins that control proper physiological development^11,12^. These examples highlighted the importance of investigating smORFs and paved the way for work in higher organisms. Recently, several mammalian microproteins have been characterized with fundamental roles in cell biology ranging from DNA repair^13^, mitochondrial function^14,15^, and RNA regulation^16^. In addition, microproteins that regulate physiological processes including muscle development^17^, muscle function^18,19^, and metabolism^20^ have been discovered. Together these studies demonstrated that genomes contain many functional smORFs and renewed interest in annotating all protein-coding smORFs.

Advances in proteomics and next-generation sequencing (NGS) technologies have provided the experimental tools necessary to identify protein-coding smORFs. The integration of RNA-Seq and proteomics approaches identified hundreds of novel microproteins in human cell lines^21,22^. Combining smORF prediction with proteomics has also uncovered more than a thousand human microproteins^23^. While proteomics provides evidence that a smORF produces a relatively stable microprotein of sufficient abundance for detection, it is limited in detecting microproteins that are not abundant or do not generate detectable peptides. With the development of ribosome profiling (Ribo-Seq), NGS can be utilized to identify ORFs that are undergoing active translation with high sensitivity and accuracy^24,25^. Ribo-Seq combines ribosome footprinting with deep sequencing to reveal the position of elongating ribosomes throughout the transcriptome^24^. Ribo-Seq has been applied successfully to smORF discovery in fruit flies^26^ and zebra fish^27^, identifying hundreds of novel translated smORFs, which is significantly more than were detectable by mass spectrometry in these organisms.

Major questions about human smORFs remain. In particular, we are interested in the following questions: (1) How many protein-coding smORFs are in the human genome? (2) Can we find evidence that smORF expression is regulated? (3) Are some smORFs ubiquitously expressed and others specific to a cell line or tissue? To answer these questions, we developed a top-down workflow that combines genome-wide transcription and translation data from RNA-Seq and Ribo-Seq, respectively, to maximize smORF discovery. Preliminary experiments revealed that smORF translation prediction by Ribo-Seq is noisier than for annotated ORFs, which led to the inclusion of reproducibility to assess the confidence of smORF annotations. We used this workflow to discover functional smORFs in HEK293T, HeLa-S3, and K562 cells, detecting over 2,500 confidently annotated protein-coding smORFs and over 7,500 in total. We also demonstrated that while smORF-encoded microproteins have distinguishing properties from annotated proteins, their expression is similarly regulated during cell stress and they are also presented as cell surface antigens. These results provide a rigorous annotation of human smORFs, dramatically increasing the coding potential of the genome, and suggest that many biologically functional microproteins are yet to be characterized.

## Results

### A Ribo-Seq and RNA-Seq based top-down smORF annotation workflow

Ribo-Seq maps the position of elongating ribosomes throughout the transcriptome through the use of cycloheximide to stall elongating ribosomes, followed by footprinting with RNase I^28^ (Fig. 1). The resulting 28-29 nt ribosome protected mRNA fragments (RPFs) are then sequenced and mapped to the transcriptome. Analyzing the global distribution and frequency of ribosomes throughout the transcriptome allows one to estimate translation levels for known ORFs and is also a highly sensitive method for identifying novel ORFs that are undergoing translation^28^. High-resolution Ribo-Seq datasets will display >70% of RPFs aligned in-frame with known coding sequences (CDS)^29,30^, captured by metagene analysis, which affords accurate identification of unannotated protein-coding smORFs.

**Figure 1.**
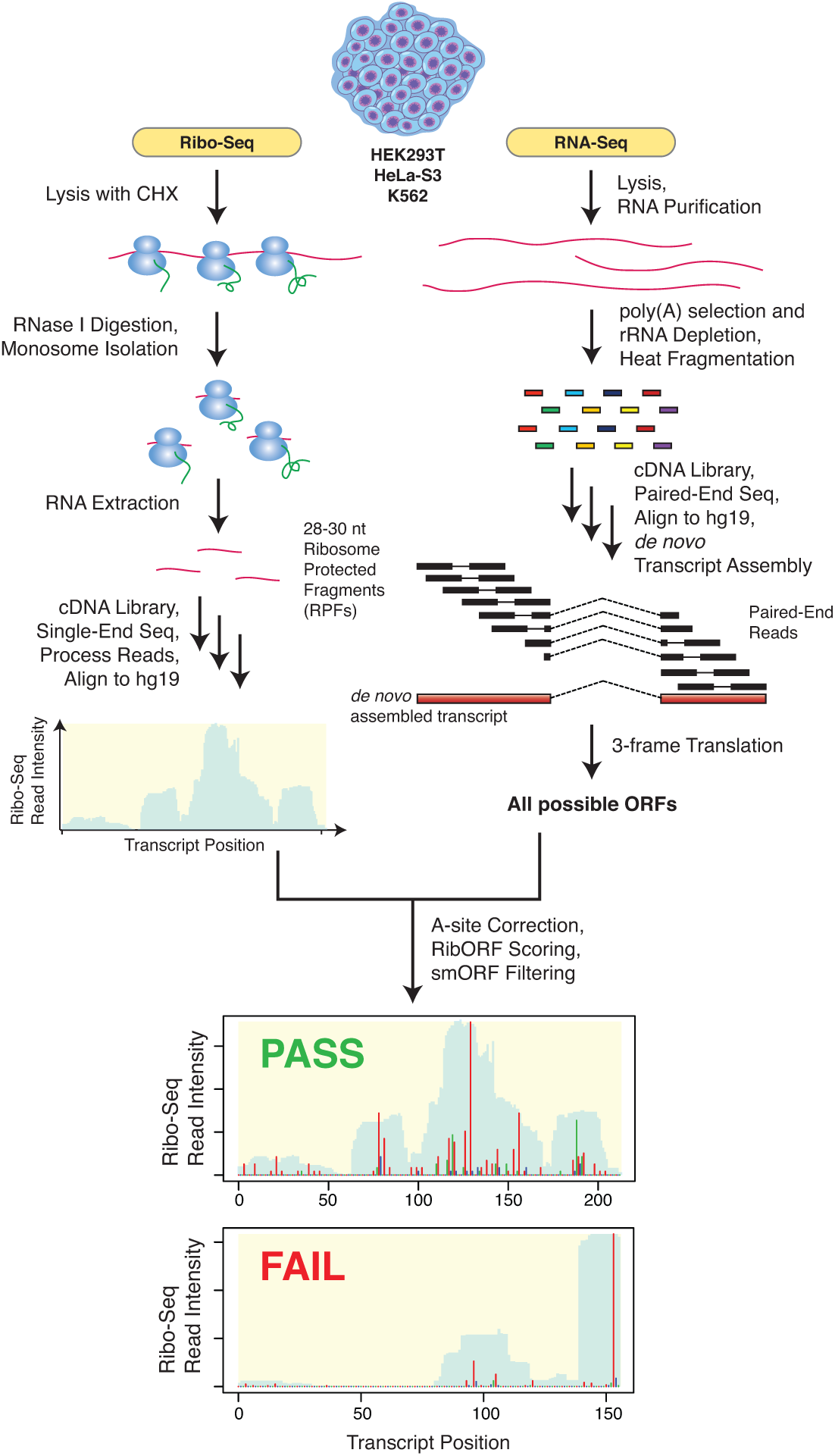
Outline of top-down smORF annotation work combining *de novo* transcriptome assembly and Ribo-Seq. RNA-Seq and Ribo-Seq datasets were collected for HEK293T, HeLa-S3, and K562 cell lines and utilized for the prediction of novel translated smORFs. For RNA-Seq, mRNA was prepared using poly-A selection and rRNA depletion to increase transcript detection. Next, a minimum of 250 million paired-end 125 base reads were collected across all cDNA libraries generated for each cell line to achieve high transcriptome coverage. RNA-Seq reads were then *de novo* assembled into a transcriptome using Cufflinks^75^. Finally, the assembled transcriptome for each cell line was *in silico* 3-frame translated to create a database of all possible ORFs. In parallel, multiple biological replicates of Ribo-Seq data were also collected for each cell line and utilized to assess translation of all smORFs in the accompanying 3-frame database. For each replicate, RibORF was used to define the A-site position of each ribosome protected fragment (RPF) and then score each smORF for translation. Those smORFs which passed RibORF scoring, did not overlap with annotated ORFs, and lacked significant similarity to RefSeq annotated proteins were retained. An example of a smORF passing RibORF scoring with coverage over the entire smORF and a high percentage of in-frame A-site reads (red) and an example of a smORF failing RibORF scoring due to poor overall smORF coverage are shown at the bottom. Predicted novel protein-coding smORFs were considered high confidence if they were found in multiple replicates.

Typically, Ribo-Seq reads are mapped onto reference transcriptome databases, such as RefSeq^31^ or Ensembl^32^, which are not representative of every cell type. Our top-down workflow maps the Ribo-Seq reads onto transcripts obtained by *de novo* assembly of RNA-Seq data obtained from the same sample. This approach identified entirely novel transcripts as well as isoforms of annotated transcripts. For example, we observed many 5’- and 3’-extensions of annotated transcripts in the *de novo* assembled transcriptome, which is important to capture given the prevalence of translated smORFs found within 5’-untranslated regions (5’-UTRs)^33–37^. We then defined ORFs across all three reading frames of the *de novo* assembled transcriptome to generate an ORF database, which includes smORFs. Incorporating *de novo* transcriptome assembly allows for more comprehensive smORF discovery in a particular sample.

After obtaining Ribo-Seq data, we scored all ORFs in the database for translation using RibORF (Fig. 1), a support vector machine-based classifier of translation^38^. RibORF uses the fraction of RPF reads aligned in-frame with the candidate ORF to calculate the overall probability of translation. This metric is dependent upon the resolution of the dataset, as higher resolution data have a greater percentage of RPF reads aligned to coding regions. RibORF also factors the uniformity and distribution of RPF reads over the entire ORF into its score. This helps filter out ORFs with high Ribo-Seq coverage in only a small region, which would likely represent an artifact^38^.

Following RibORF scoring, the list of predicted translated ORFs was filtered by size to remove all ORFs smaller than 6 codons, which are not amenable to detection by tandem mass spectrometry, and greater than 150 codons, because protein-coding smORFs larger than the usual cutoff of 100 codons have been reported^21,39-41^. Next, translated smORFs found to overlap with annotated CDS regions in the UCSC database were removed. This step filtered out several thousand smORFs comprising both annotated genes as well as putative smORFs that overlap out-of-frame with an annotated gene, which can be difficult to accurately score. Finally, this set of smORF-encoded microproteins was analyzed for similarity to human RefSeq proteins by BLASTp. Only low scoring hits were retained, which removed pseudogenes and any missed annotated genes. The remaining hits constitute the set of novel microprotein-encoding smORFs (Fig. 1).

### Annotating protein-coding smORFs in HEK293T cells

We tested the top-down smORF annotation workflow in HEK293T cells because we previously identified dozens of microproteins in these cells by proteomics^21^. Ribosome footprints from HEK293T cells were prepared using a protocol that afforded sub-codon resolution with HEK293 cells^30^. Initially, only ~50% of reads aligned in-frame by metagene analysis, and RPF lengths peaked at 31 nt (Fig. 2a). While this resolution is comparable to several published datasets (Supplementary Fig. 1), we endeavored to collect higher resolution datasets as well to ensure identification of translated smORFs that require higher accuracy read alignment. The 31 nt peak footprint length indicated that the nuclease digestion step, which serves to trim away all unprotected RNA, was incomplete. To gain finer control over nuclease digestion, we followed a reported strategy that normalizes the amount of nuclease added to the RNA concentration^29^, as opposed to adding nuclease based on a cell confluency or the lysate’s A260 value. We were able to generate two additional HEK293T Ribo-Seq datasets with ~60% and >70% of reads in-frame by metagene analysis and RPF lengths that peaked at 30 and 28 nt, respectively (Fig. 2a). Given that published datasets show a wide range of resolutions (Supplementary Fig. 1), we carried all three datasets, low-, medium-, and high-resolution, forward for protein-coding smORF prediction.

**Figure 2.**
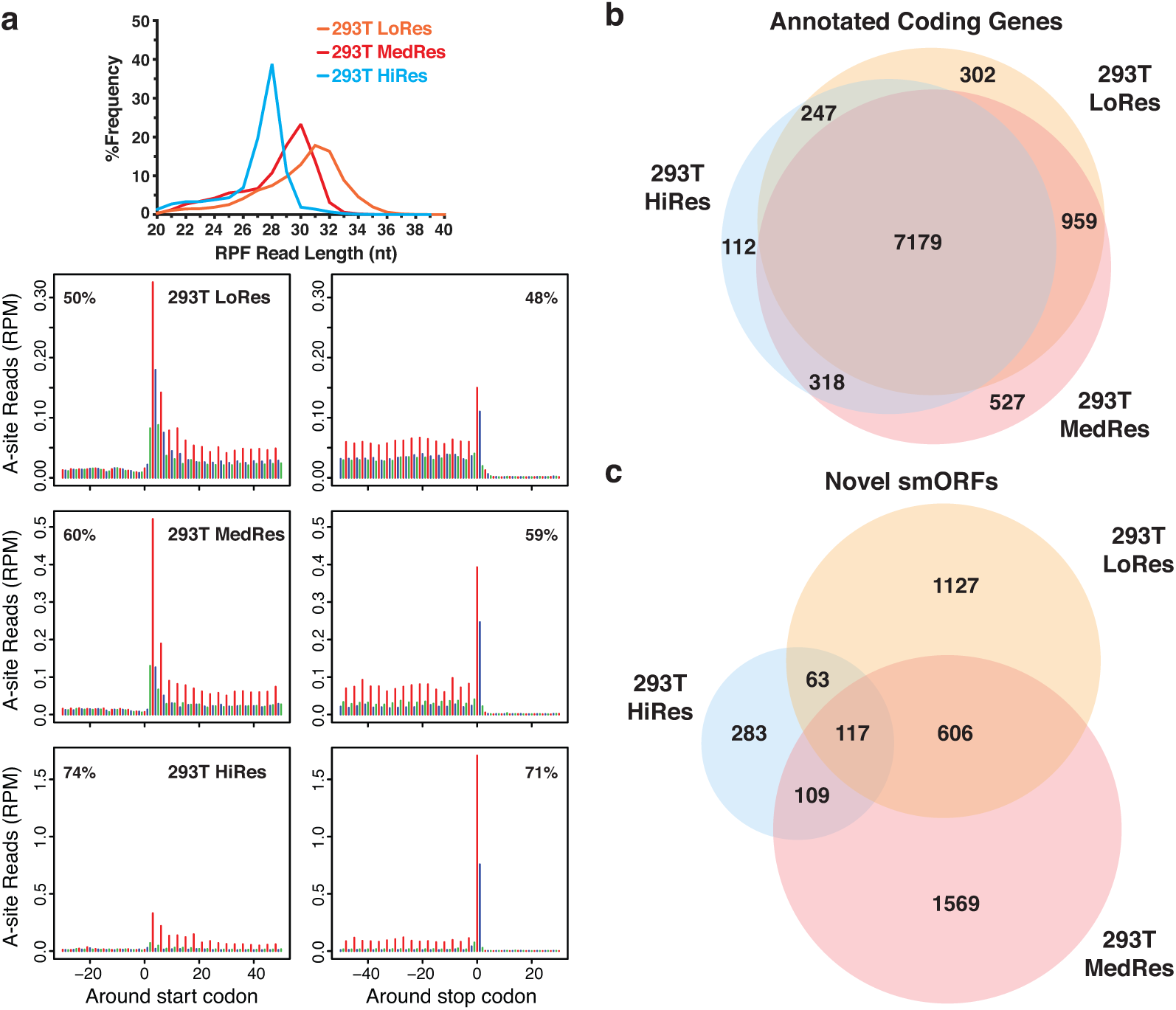
Number of novel protein-coding smORFs predicted varies with overall Ribo-Seq resolution. **a** Three replicates of HEK293T samples were subjected to increasing RNase I nuclease digestion resulting in a range of overall Ribo-Seq resolutions: low (LoRes), medium (MedRes), and high (HiRes). **a, top** RPF read length distribution plot showing the differences in read size frequencies across replicates. The expected ribosome footprint size is 28 nt. **a, bottom** Metagene plots showing RPF read alignment around the start site and stop site for each 293T replicate created using RibORF. The 5’-position of each RPF read was shifted to the ribosomal A-site and then mapped to all hg19 RefSeq coding transcripts, which were used to construct the metagene. The metagene coding region is aligned to frame 1 (red), while frame 2 (blue) and frame 3 (green) are out of frame. The percentage of reads aligned to the coding region is noted in the top corner. Higher percentages equate to higher resolution datasets. 28-34 nt reads were used for the LoRes metagene plot, 29-33 nt for MedRes, and 25-29 nt for HiRes. **b** Venn diagram showing overlap of annotated RefSeq genes passing RibORF scoring between all three HEK293T Ribo-Seq replicates. **c** Venn diagram showing overlap of novel protein-coding smORFs passing RibORF scoring and our smORF filters between the three replicates.

Several previous studies have combined multiple Ribo-Seq experiments, even across different cell lines, to maximize the available reads and increase the likelihood of passing translation scoring^29,30,38,42^. However, this strategy also allows for more false positives when the same thresholds are applied due to reads accumulating on a candidate ORF either because of non-productive ribosomal binding or noise inherent to the Ribo-Seq protocol^43^. In addition, combining experiments does not allow one to assess the reproducibility of translation predictions. Indeed, Ribo-Seq replicates can show high correlation gene counts but still have low local positional correlation^43^, which affects translation scoring of novel ORFs. Furthermore, validating results using replicates has been shown to be critical in other NGS-based assays^44^. If multiple experiments are analyzed separately, novel smORFs predicted as translated in every experiment regardless of noise and sequencing depth are more confidently protein-coding than those found in a single experiment. This allows one to also observe how differences in RPF preparation affect translation scoring. Therefore, we analyzed each Ribo-Seq experiment for smORF translation separately.

We first used RibORF to score translation of canonical RefSeq genes to confirm the quality of our Ribo-Seq datasets and determine the noise level for *bona fide* genes. Despite differences in sequencing depth and resolution, we observed high overlap among the 9,644 canonical genes called translated, with 74% found in all three experiments (Fig. 2b). These results demonstrated the quality of our datasets and suggested that their resolution is sufficient to predict translation. For smORFs, however, we found that resolution has a strong influence on the total number of translated smORFs and that translation prediction is much noisier (Fig. 2c). Utilizing our smORF annotation workflow, we were able to identify 1,913 (low-resolution), 2,401 (medium-resolution), and 572 (high-resolution) novel translated smORFs (Fig. 2c), totaling 3,874 unique hits in all. Of these, 117 smORFs were scored as translated in all three experiments and 895 smORFs in at least two experiments (Fig. 2c). Despite the low overlap, taking only the set of smORFs found in multiple experiments dramatically increases the number of protein-coding smORFs detected in previous proteomics-based efforts^21,22,39,45^. We were also able to validate translation of several smORFs identified previously by proteomics in HEK293T cells^21^. Interestingly, 606 smORFs were found in both lower resolution datasets but not the high-resolution dataset, suggesting that there is a benefit to collecting datasets of varying resolution. These results support the idea that our combined Ribo-Seq and RNA-Seq workflow is an effective and sensitive means for identifying novel translated smORFs. Practically, these data also indicate that we must run several Ribo-Seq experiments for each sample to more confidently annotate smORFs.

### Regulation of smORF transcription and translation during endoplasmic reticulum stress

Having identified thousands of novel protein-coding smORFs in HEK293T cells, we next searched for evidence of regulation as a means to uncover possible biological roles, and chose to look at changes in transcription and translation induced by endoplasmic reticulum (ER) stress. ER stress leads to the accumulation of unfolded or mis-folded proteins and triggers a well-characterized signaling cascade dubbed the Unfolded Protein Response (UPR)^46^. The UPR attempts to restore ER homeostasis if possible and promote survival, or induces apoptosis if the stress is unresolvable. Thapsigargin (TG) and tunicamycin (TM) are small molecules that can trigger ER stress. TG inhibits the ER calcium pump SERCA, causing reduced calcium levels in the ER and inhibition of calcium-dependent chaperones, while TM inhibits the ER glycoprotein transferase, which prevents sugar conjugation to proteins and causes protein mis-folding.

HEK293T cells were treated with TG or TM for 4 h and compared to vehicle-treated cells. RNA-Seq and high resolution Ribo-Seq data were collected for each sample to monitor changes in mRNA expression and translation (Supplementary Fig. 3a and 4), and to identify any additional novel protein-coding smORFs. Applying our smORF annotation workflow, we identified an additional 666 smORFs, increasing the total count of unique translated smORFs to 4,540 across all HEK293T datasets collected (Supplementary Data 2). Confirming activation of the UPR by TG and TM treatment, HSPA5, HYOU1, DDIT3, and other known UPR genes were upregulated^47^ (Fig. 3a and Supplementary Data 1). Gene Ontology (GO) analysis of TG- and TM-regulated genes also confirmed activation of the UPR (Supplementary Data 1).

**Figure 3.**
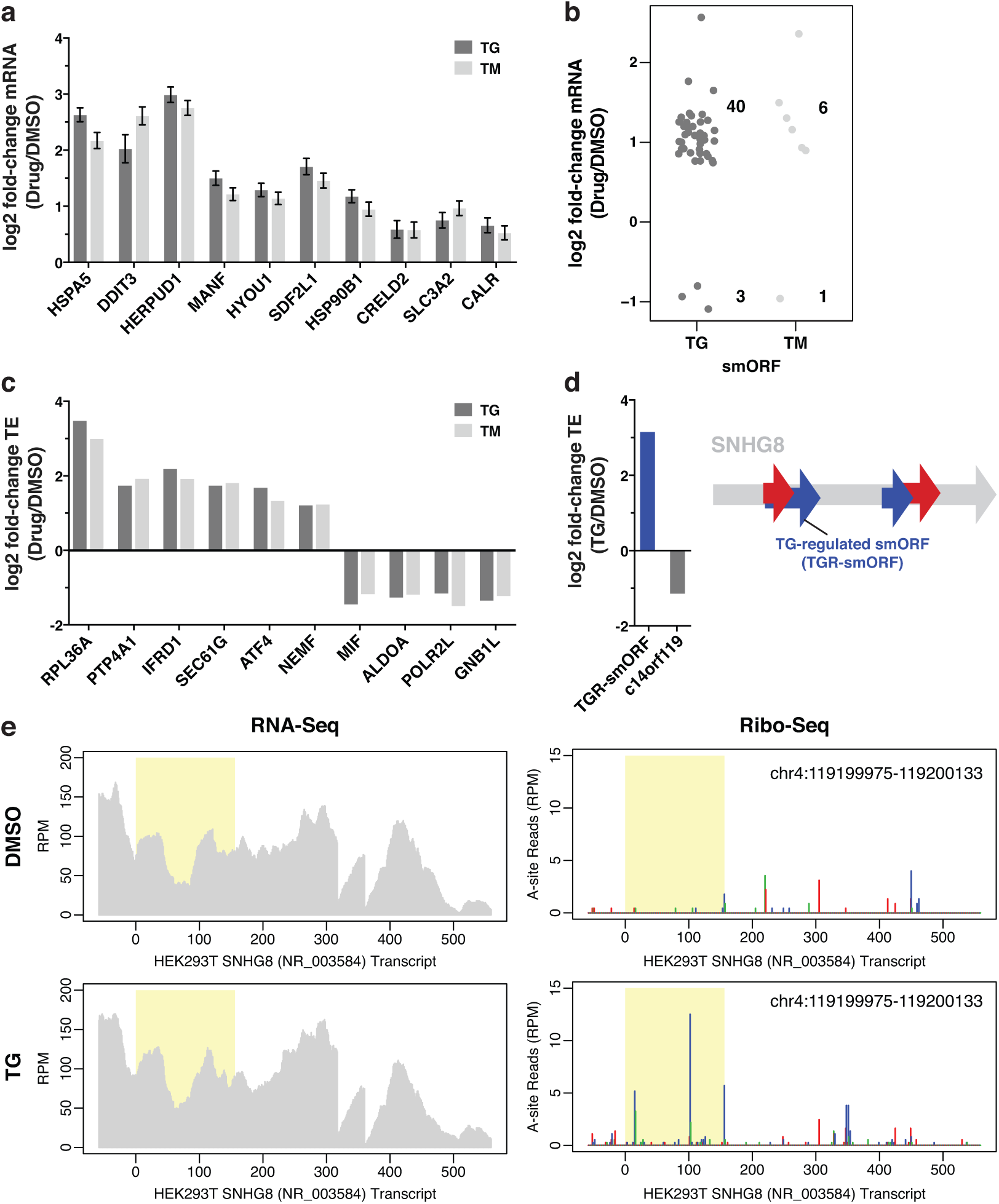
smORF expression is regulated during endoplasmic reticulum (ER) stress. **a** Bar graph showing log2 normalized fold-change in expression of canonical ER stress response genes after 4 h treatment of HEK293T cells with 1 µM thapsigargin (TG) or 5 µg/mL tunicamycin (TM) relative to DMSO as measured by RNA-Seq (error bars represent the log fold change SE, p_adj_ < 0.05). Two biological replicates for each condition were analyzed. **b** Strip chart showing the change in expression of novel smORFs induced by TG and TM (p_adj_ < 0.05). Only smORFs identified in at least two Ribo-Seq experiments across all HEK293T replicates were considered. **c** Bar graph showing log2 normalized fold-change in translation efficiency (TE) of canonical ER stress response genes after treatment with TG or TM as calculated by Xtail50 (padj < 0.1). **d, left** Bar graph showing the change in TE induced by TG of a novel smORF on the non-coding RNA SNHG8 and an annotated smORF, c14orf119. **d, right** Schematic showing the SNHG8 transcript and the novel smORF that is translationally regulated in response to TG, TGR-smORF. Four novel smORFs were identified at least twice on SNHG8: two in frame 1 (red) and two in frame 2 (blue). **e** Representative RNA-Seq read coverage and ribosomal A-site plots (Ribo-Seq) for SNHG8 showing the change in ribosome occupancy induced by TG. TGR-smORF is highlighted by the yellow box and is found frame 2 (blue). TGR-smORF coordinates are shown in the top corner. The y-axis shows the intensity of read peaks in reads per million (RPM).

Next, we analyzed the novel smORFs for transcriptional regulation under ER stress, focusing on the 1,409 smORFs identified in at least two Ribo-Seq experiments (Supplementary Data 2). TG and TM induced significant changes in the mRNA expression of 43 and 7 smORFs, respectively (Fig. 3b). This suggests that the encoded microproteins might function as part of the UPR. In addition, many of these smORFs were found within the 5’-UTR of a known protein-coding gene, termed upstream ORFs (uORFs) (Supplementary Data 2), which are frequently found on critical UPR genes, such as ATF4, DDIT3, IBTK, and GADD34^48,49^. Interestingly, transcriptionally regulated smORFs were also found on annotated non-coding RNAs (ncRNAs), including multiple small nucleolar RNA host gene (SNHG) family members. Thus, these ncRNAs may have dual roles in the UPR as functional RNAs and by encoding microproteins.

Monitoring transcriptional changes is helpful in identifying functionally relevant genes, however, if a gene is translationally regulated then changes in mRNA levels will not correlate with changes in protein level. Indeed, several UPR pathway genes are translationally regulated during ER stress^40,41,48,49^. To account for this possibility, we also analyzed genes for differential translation using Xtail^50^. Xtail uses Ribo-Seq data to measure changes in translation while accounting for changes in transcript abundance using RNA-Seq data to quantify the magnitude and statistical significance of differential translation efficiency (TE) genome-wide. Both TG and TM induced higher TE for ATF4, IFRD1, and SEC61G, which have been shown to be regulated during ER stress^51–53^, as well as several other genes (Fig. 3c). In total, 129 annotated genes had significantly regulated TEs in response to TG treatment, while 25 genes were regulated in response to TM treatment (Supplementary Fig. 2 and Supplementary Data 1). GO analysis of the 129 TG-regulated genes showed enrichment for RNA processing and splicing (Supplementary Data 1).

Analysis of HEK293T smORFs revealed a robust increase in the translation efficiency of a smORF within SNHG8, which we named thapsigargin-regulated smORF (TGR-smORF) (Fig. 3d). This change is clearly visualized by comparing the Ribo-Seq read coverage plots for SNHG8 (Fig. 3e). While there was no significant change in SNHG8 transcript levels between vehicle- and TG-treated cells (Fig. 3e, left), the Ribo-Seq read plot reveals a dramatic increase in the ribosome occupancy within TGR-smORF (Fig. 3e, right). In addition, the TE of the annotated but uncharacterized smORF c14orf119 significantly decreased in response to TG (Fig. 3d). Thus, TGR-smORF and c14orf119 are candidates for further functional studies to determine what role they serve in UPR. Together, these ER stress-regulated smORFs demonstrate the value in using genomics to identify smORFs associated with a particular biology.

### Annotation of protein-coding smORFs in additional human cell lines

Our smORF annotation workflow uncovered more than 4,000 novel protein-coding smORFs across nine Ribo-Seq datasets in HEK293T cells. Next, we wanted to determine whether this number is specific to HEK293T or if other cell lines would provide similar results. Furthermore, analysis of additional cell lines would enable us to determine if there are smORFs with cell line-specific or more ubiquitous expression. Because they differ in their tissue of origin from HEK293T, we selected the cell lines K562, chronic myeloid leukemia derived, and HeLa-S3, cervical cancer derived. As ENCODE cell lines, these also provide a wealth of high quality genomic, transcriptomic, and functional data available for follow-up analyses^54^.

As with HEK293T, HeLa-S3 cell lysates were digested using different conditions to maximize the number and accuracy of smORFs identified. Metagene analysis showed a range of resolutions across the four datasets collected, from ~50-70% reads in-frame (Supplementary Fig. 3b and 5). Altogether, 2,614 novel smORFs were scored as translated, with 777 smORFs found in at least two experiments (Supplementary Data 2). The overall resolution and RPF length distributions for the HeLa-S3 datasets were similar to the HEK293T datasets collected using similar digestion conditions (Fig. 2a, Supplementary Fig. 5). Next, we collected three Ribo-Seq datasets from K562 using a range of digestion conditions. Because K562 cells were grown in suspension, they were subjected to longer periods of CHX treatment during washing. This caused enrichment in start site reads (Supplementary Fig. 6), as was observed in previous studies^55,56^. Despite the longer CHX treatment, 2,464 novel protein-coding smORFs were identified in K562 cells, with 542 smORFs found in at least two experiments (Supplementary Data 2). All conditions tested resulted in >75% reads in-frame by metagene analysis and RPF length distribution peaking at 28-nt (Supplementary Fig. 3c). However, K562 HiRes3 displayed a broader footprint length distribution (Supplementary Fig. 6).

**Figure 5.**
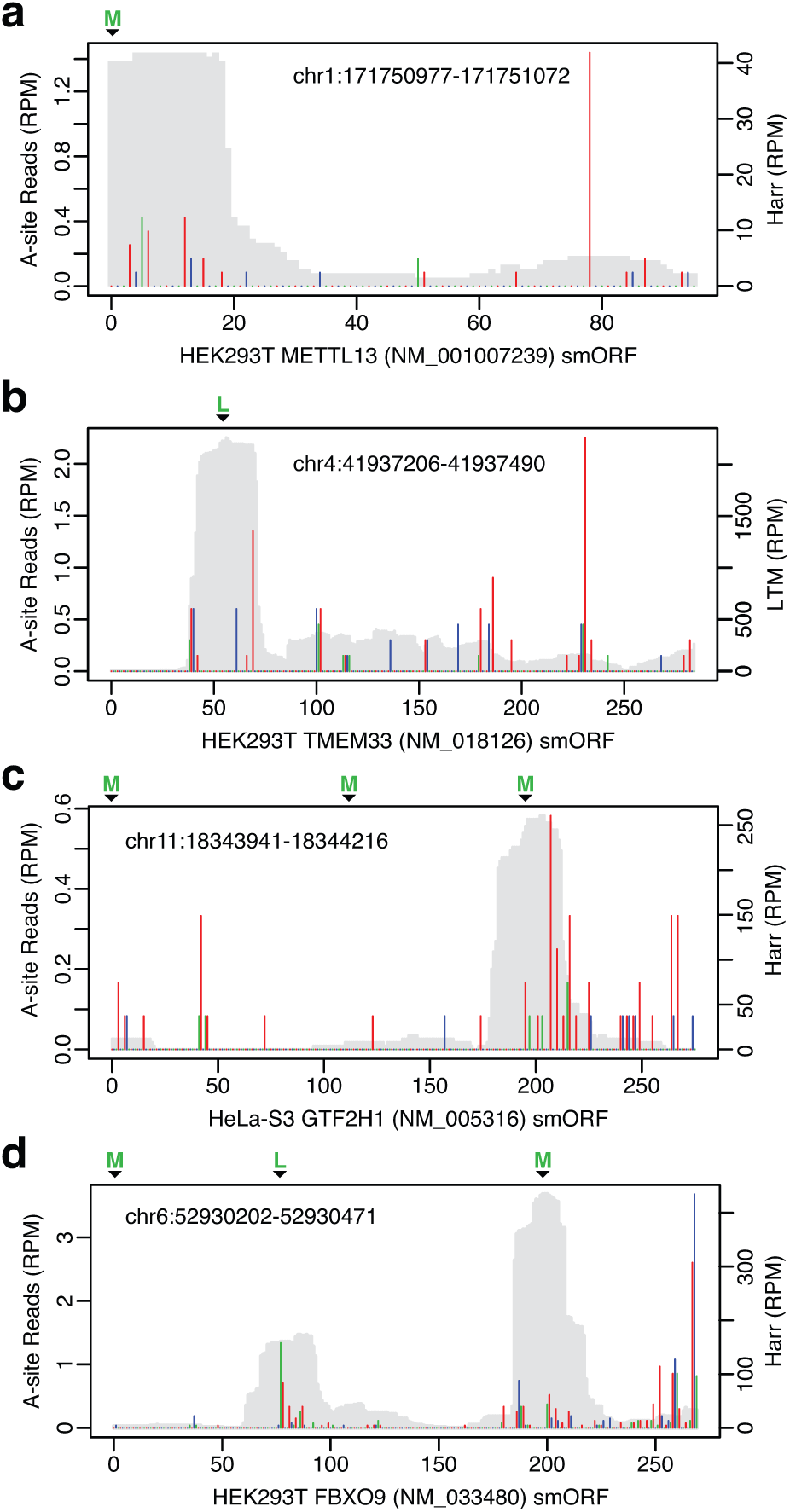
Translation inhibitors lactimidomycin (LTM) and harringtonine (Harr) aid in identifying smORF start sites. **a** Read coverage peak in Harr-treated cells confirmed translation initiation at the upstream-most AUG start codon (encoding methionine, denoted by the green M) in the novel protein-coding smORF occurring within the 5’-UTR of methyltransferase like 3 (METTL3). Elongating ribosomal A-site reads are depicted as bars and are color coded by reading frame. The smORF coding region is aligned with frame 1 (red). RPF read coverage in Harr-treated cells is shown in grey. **b** RPF read coverage in LTM-treated cells identified the start site as occurring at the near-cognate start codon UUG (encoding leucine, denoted by green L) in the non-AUG smORF occurring within the 5’-UTR of transmembrane protein 33 (TMEM33). **c** RPF read coverage in Harr-treated cells identified a downstream AUG start codon (denoted by third green M) as the predominant translation initiation site in the smORF occurring within the 5’-UTR of general transcription factor IIH subunit 1 (GTF2H1). A-site coverage upstream of the third AUG codon and a small Harr peak suggested that the first AUG may also function as the start site to a lesser extent. **d** RPF Read coverage in Harr-treated cells showed mixed start site usage between a downstream near-cognate start codon CUG (encoding leucine, denoted by green L) and a downstream AUG start codon (encoding methionine, denoted by second green M) in the smORF occurring within an alternative 5’-UTR of F-box protein 9 (FBXO9). A-site coverage supports translation from both start sites.

**Figure 6.**
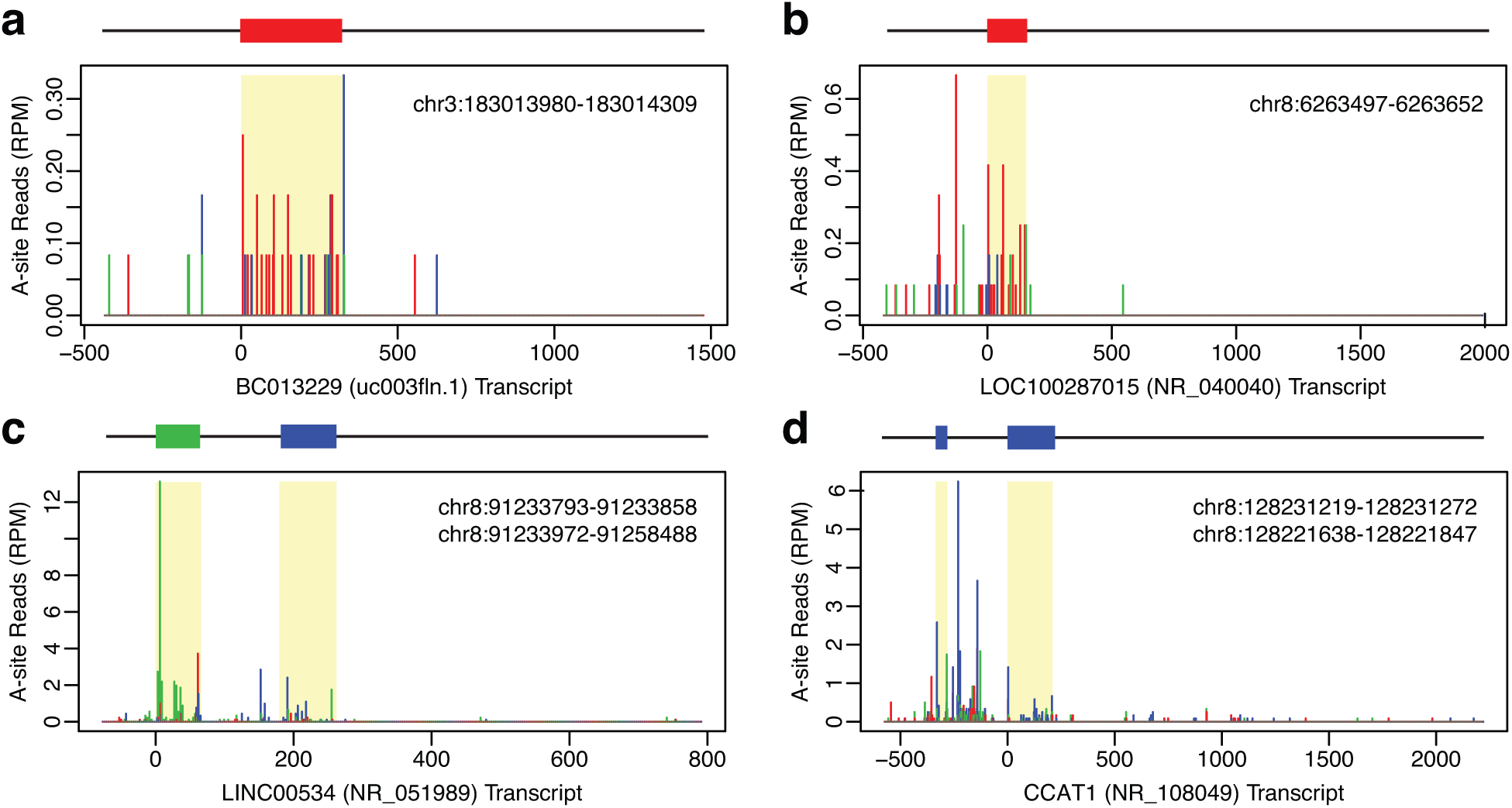
Ribo-Seq identified hundreds of novel protein-coding smORFs on annotated non-coding RNAs (ncRNAs). Ribo-Seq identified hundreds of smORFs on RefSeq and UCSC annotated non-coding RNAs across all three cell lines. **a-b** The UCSC ncRNA BC013229 and the RefSeq ncRNA LOC100287015 each contain a novel protein-coding smORF in frame 1 (red) that was identified in all HeLa-S3 Ribo-Seq experiments. The smORF coordinates are shown in the top corner. Both ncRNAs are currently uncharacterized. **c** The uncharacterized RefSeq ncRNA LINC00534 is polycistronic, containing two confident novel protein-coding smORFs, one in frame 3 (green) and one in frame 2 (blue). Both smORFs were identified in all K562 Ribo-Seq experiments. **d** The ncRNA colon cancer associated transcript 1 (CCAT1) is polycistronic with at least two confident novel smORFs, both in frame 2 (blue). The upstream smORF was identified in two out of four HeLa-S3 Ribo-Seq experiments, and the larger downstream smORF passed in all HeLa-S3 experiments.

Between the three cell lines profiled, we identified 7,554 novel protein-coding smORFs across three diverse tissue types (Supplementary Data 2). Most smORFs are only identified in a single experiment, but there are many smORFs that overlap between cell lines or are found in multiple experiments from a single cell line. In total, 483 smORFs were detected in all three cell lines, 1,581 in at least two cell lines, and 2,689 in at least two experiments (Fig. 4a). We also observed that smORFs detectable in two or more cell lines are more likely to utilize AUG as an initiation codon than smORFs found in only one cell line. These results reveal that smORFs, like annotated genes, can be ubiquitous and cell type specific.

**Figure 4.**
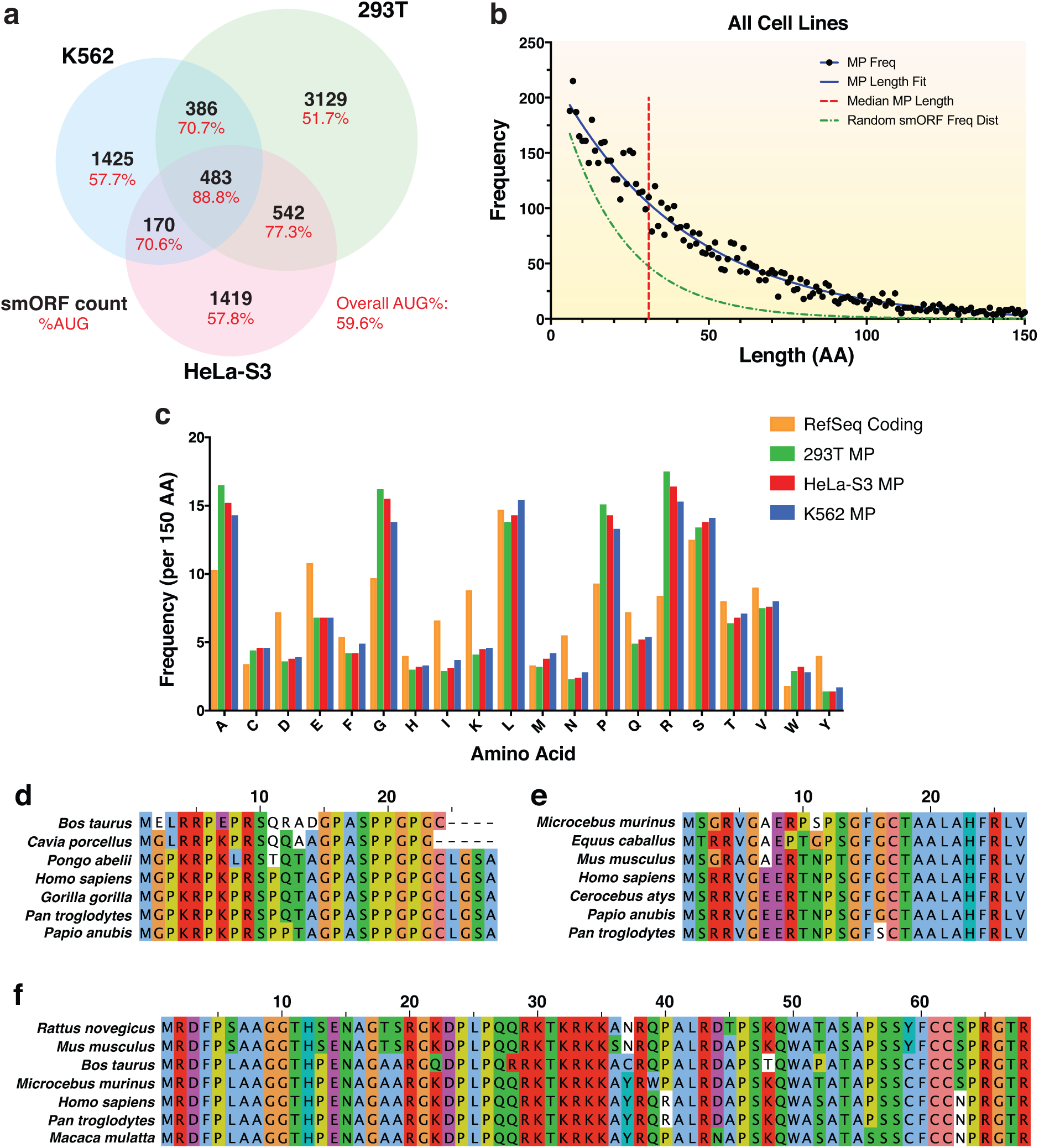
Characteristics of novel protein-coding smORFs identified across three different human cell lines and conservation in other mammalian species. **a** Venn diagram showing the overlap of the 7,554 novel smORFs identified in HEK293T, HeLa-S3, and K562 cell lines (black text). The percent of smORFs containing an AUG start codon for each sector is also shown (red text). **b** Scatter plot showing the frequency distribution of smORF-encoded microprotein (MP) lengths in amino acids (aa). The median microprotein size is 32 aa (red). The MP length distribution can be fit with a decay curve of the formula N_0_e-λx, where N_0_ = 224 and λ = 0.024 (blue). This is a slower decay than the expected frequency distribution of randomly occurring MPs based on the probability of encountering a stop codon, where λ = 0.05 (green)^8^. **c** Frequency of aa occurrence per 150 aa for annotated RefSeq proteins and novel microproteins identified in each cell line. **d** Sequence alignment for a novel microprotein encoded by a smORF within the 5’-UTR of four jointed box 1 (FJX1). FJX1 smORF has an average PhyloCSF score of 3.49 using the 29-mammal alignment, indicating a higher probability of being protein-coding. **e** Sequence alignment for a novel microprotein encoded by a smORF within the 5’-UTR of nuclear casein kinase and cyclin dependent kinase substrate 1 (NUCKS1). This smORF has a negative PhyloCSF score, but the microprotein sequence shows high similarity to translated regions in mammalian species by tBLASTn. **f** Sequence alignment for a novel microprotein encoded by a smORF within the 5’-UTR of B-cell CLL/lymphoma 9 (BCL9). This smORF has a negative PhyloCSF score, but the microprotein sequence shows high similarity to proteins in other mammalian species by BLASTp and tBLASTn.

### Analyzing microprotein properties and sequence conservation across species

Next, we analyzed microprotein properties for comparison to annotated proteins. The median length of the encoded microproteins is 32 amino acids (Fig. 4b, red line), whereas the median human protein length in the Pfam database is 416 amino acids^57^. The frequency distribution of microprotein lengths can be fit by the decay formula N_0_e^−λx^ where N_0_ = 224 and λ = 0.024 (Fig. 4, blue line), which has a slower decay than the expected frequency distribution of microprotein length based on the random occurrence of a stop codon after an initiation codon^8^ (Fig. 4b, green line). Thus, protein-coding smORFs occur at a higher frequency than expected by chance.

Each cell line was then analyzed independently to assess the overall amino acid composition of smORF encoded microproteins. The data revealed a clear difference in several amino acid frequencies that distinguish microproteins from annotated RefSeq proteins (Fig. 4c). The amino acids with markedly higher frequencies include alanine, glycine, proline, and arginine, while cysteine and tryptophan were slightly increased. Several amino acids including aspartic acid, glutamic acid, isoleucine, lysine, asparagine, glutamine, and tyrosine occur less frequently. Interestingly, the amino acid composition of microproteins are similar across all three cell lines.

We also searched for structural features, including transmembrane helices and conserved protein domains, to understand how microproteins compare to annotated proteins. Given the inherently small size of microproteins (Fig. 4b), we did not anticipate many to contain canonical structural motifs. Using TMHMM2.0^58^, we only identified 48 microproteins found in at least two Ribo-Seq experiments (1.8%) with predicted transmembrane helix domains (Supplementary Data 3). In addition, only 17 microproteins (0.06%) contain known protein domains based on the Conserved Domain Database^59^. These results are consistent with most microproteins employing different structures from annotated proteins.

Despite these differences, we hypothesized that many microproteins would show sequence conservation across other mammalian species, similar to annotated proteins. We first employed PhyloCSF, which uses a multi-species nucleotide alignment to examine sequences for signatures of conserved coding regions^60^. At least one exon with a positive average PhyloCSF score was found in 423 smORFs (Supplementary Data 2), such as the novel smORF within the 5’-UTR of *FJX1* (Fig. 4d). We also searched for sequence similarities across other species using tBLASTn and BLASTp as evidence for possible protein conservation. Using tBLASTn, 4,687 microproteins were found to have high similarity to translated RNA sequences from at least one other species, including 273 to mouse sequences (Supplementary Data 2). Additionally, 476 microproteins with high similarity to known and predicted proteins were found in other species using BLASTp (Supplementary Data 2). In many instances, clear sequence similarity was observed across several species using tBLASTn and BLASTp despite having negative PhyloCSF scores (Fig. 4e,f). These data demonstrate that a large portion of our novel microproteins are likely conserved.

### Identifying smORF Translation Initiation Sites

Approximately 40% of the predicted protein-coding smORFs lack an in-frame canonical AUG start codon (Fig. 5a), making their translation initiation sites difficult to identify. Fortunately, one of the most powerful features of Ribo-Seq is the ability to empirically identify translation start sites through treatment with initiation-specific inhibitors, such as harringtonine (Harr) and lactimidomycin (LTM)^25,61,62^. For example, Harr treatment induced RPF accumulation centered on the first AUG start codon in a novel *METTL3* uORF (Fig. 5a). Start site inhibitors also proved capable of identifying alternative initiation codons. LTM treatment enriched RPF coverage over the near cognate start codon UUG in a *TMEM33* uORF (Fig. 5b), supporting its translation despite the lack of an in-frame AUG start codon.

These inhibitors were also helpful in identifying the predominant codons for translation initiation when multiple canonical or near cognate start codons were present. For example, there are three in-frame AUG codons within a novel uORF on *GTF2H1*. Surprisingly, Harr treatment induced the highest RPF accumulation over the third AUG codon, with only a small peak present over the first AUG (Fig. 5c), suggesting that both a long and predominant short form of the microprotein are made. Similarly, we saw mixed start site usage for the uORF on *FBXO9* (Fig. 5d). We observed translation initiation peaks over a CUG codon and a downstream AUG codon, suggesting that both are also utilized to produce a long and short form of the microprotein. Of note, no initiation peak was observed over the most upstream in-frame AUG codon. The ability to empirically detect initiation sites by Ribo-Seq provides invaluable information for accurately annotating smORFs and is a significant advantage over other methods.

### Novel protein-coding smORFs are found on annotated and unannotated transcripts

Having found thousands of novel protein-coding smORFs, we subsequently determined their position relative to the annotated RefSeq transcriptome. In doing so, we hoped to see how many annotated transcripts harbor translated smORFs and where within transcripts they occur most frequently. Over half of all predicted translated smORFs overlapped with RefSeq transcripts. The majority were found within the 5’-UTR of known genes (Supplementary Fig. 6), while only a small portion of smORFs were found within the 3’-UTR and on the strand opposite annotated genes. Interestingly, 623 novel smORFs were discovered on RefSeq ncRNAs, and several more on UCSC ncRNAs. Many of these smORFs are high confidence identifications found in several experiments. For instance, we found translated smORFs on the ncRNAs, *BC013229* and *LOC100287015* (Fig. 6a,b), which were identified in every HeLa-S3 experiment. We also discovered ncRNAs containing multiple protein-coding smORFs, such as *LINC00534*, which contains two novel smORFs in different reading frames (Fig. 6c). In addition, we found two confident smORFs on *CCAT1*, and several more that were called translated in only a single experiment (Fig. 6d and Supplementary Data 2). These data suggest that some ncRNAs are actually an overlooked source of coding potential in the genome.

A large portion of novel protein-coding smORFs were also located on unannotated transcripts, or those that are present in the *de novo* transcriptome assembly but not RefSeq. These unannotated transcripts comprise isoforms of annotated transcripts, containing either extensions of exons or novel exons, as well as entirely new transcripts. One *de novo* assembled transcript includes a 5’-extension of *c6orf62* which contains a translated smORF (Fig. 7a). Other examples include novel exons, such as the smORF-containing *EYA4* isoform found specifically in HeLa-S3 samples (Fig. 7b) and the *GGPS1* isoform with an alternative 5’-UTR containing a novel smORF (Fig. 7c). Furthermore, we were able to identify several protein-coding smORFs on transcripts that do not overlap with any annotated gene, and many of these uannotated transcripts are cell type specific (Fig. 8a-c). These unannotated smORFs emphasize the importance of utilizing *de novo* transcriptome assembly with Ribo-Seq to identify novel protein-coding smORFs.

**Figure 7.**
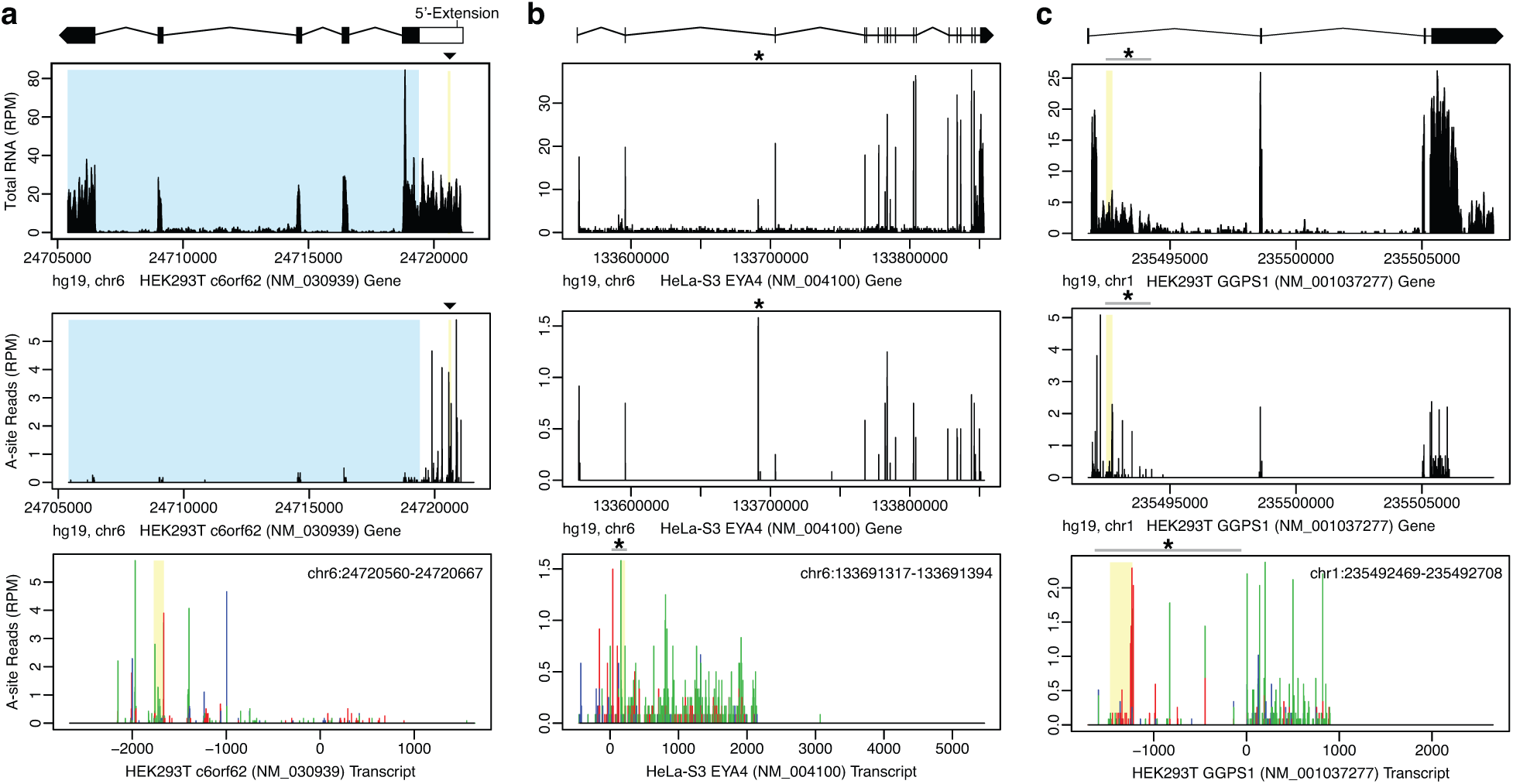
Ribo-Seq identified smORFs on unannotated *de novo* assembled transcript isoforms. **a** A novel protein-coding smORF was identified within a 5’-extension of c6orf62. The top plot shows RNA-Seq read coverage at the genomic level. The RefSeq annotated transcript is highlighted by the blue box and the novel smORF is highlighted by the yellow box and black triangle (▼) above. The exon positions are depicted by the transcript model above the plot. Black boxes represent the annotated exons, the white box represents the 5’-extension, introns are depicted by the connecting lines, and the strand orientation is noted by the arrowhead. The middle A-site plot shows Ribo-Seq coverage at the genomic level. The bottom A-site plot shows Ribo-Seq coverage at the transcript level with reads colored by frame. The novel smORF occurs in frame 3 (green). Position 0 marks the start of the annotated coding region. The smORF coordinates are shown in the top corner. **b** A novel protein-coding smORF was identified within an unannotated exon in the middle of the annotated EYA transcriptional coactivator and phosphatase 4 (EYA4) coding region, and is specific to HeLa-S3. The RNA coverage plot shows the novel exon occurring upstream of the third exon in the annotated transcript, denoted by the asterisk (**\***) and grey bar. The gene level A-site plot shows high coverage over this novel exon, and the transcript level A-site plot shows the smORF in frame 3 (green). The novel exon was assembled as the first exon in an isoform of EYA4. **c** A novel protein-coding smORF was identified within an unannotated exon upstream of the geranylgeranyl pyrophosphate synthase 1 (GGPS1) coding region. The gene level RNA and A-site Ribo-Seq plots show significant read coverage over the novel exon. The transcript level A-site plot shows the smORF in frame 1 (red). The novel exon was assembled as the first exon in an isoform of GGPS1, altering the 5’-UTR.

**Figure 8.**
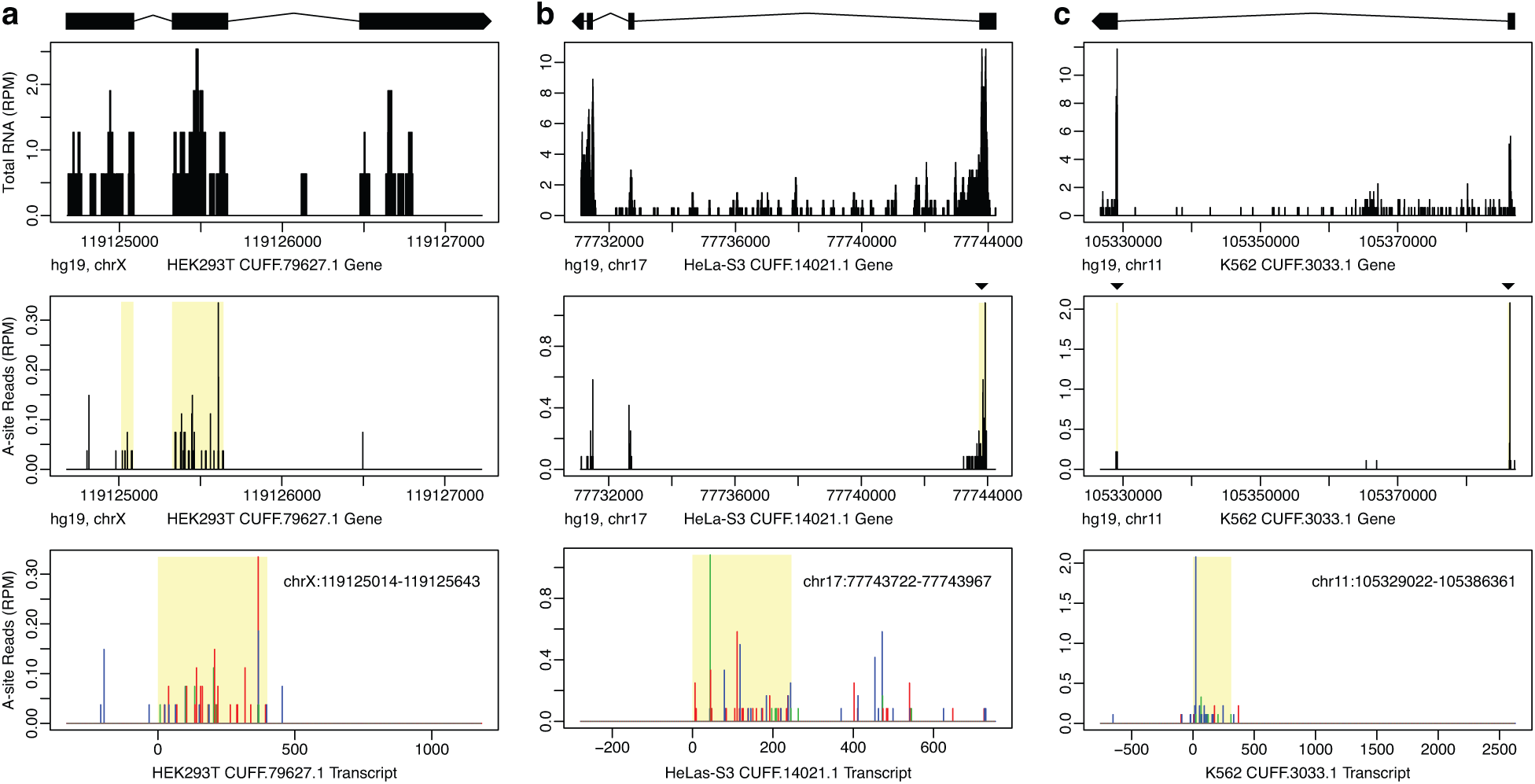
Ribo-Seq identified smORFs on novel unannotated transcripts that are also cell type specific. Novel protein-coding smORFs were identified on unannotated *de novo* assembled transcripts which had no overlap with annotated genes. Examples are shown which are specific to **a** HEK293T, **b** HeLa-S3, and **c** K562. The top plot shows RNA coverage at the genomic level with the exon model of the Cufflinks assembled transcript shown above. Black boxes represent the exons, connecting lines represent the introns, and the strand orientation is noted by the arrowhead. The middle A-site plot shows the Ribo-Seq coverage at the gene level with the smORF highlighted by the yellow box and black triangle (▼) above. The bottom A-site plot shows the Ribo-Seq coverage at the transcript level with reads colored by frame. The smORF coordinates are shown in the top corner. The smORFs in **a** and **b** are in frame 1 (red), while the smORF in **c** is in frame 2 (blue).

### Detection of novel microproteins in Human Leukocyte Antigen class I (HLA-I) peptidomics datasets

Translation of thousands of smORFs were detected by Ribo-Seq. However, Ribo-Seq cannot determine whether the encoded microprotein is sufficiently long-lived to be functional. Mass spectrometry can detect proteins that accumulate to a steady state concentration above the limit of detection, offering important complementary data. Proteins are often identified from mass spectrometry data using a search strategy that matches ms2 spectra with tryptic peptides from a protein database. However, these novel microproteins are not included in human proteome databases, and therefore would not be identified in published proteomics studies. Another challenge with most proteomics datasets is that they do not enrich for smaller peptides or small proteins prior to analysis, which we have found to be critical for microprotein detection^21,39^. Therefore, we searched published datasets that had an enrichment step built in to see if microproteins are detected when their sequences are appended to a human proteome database.

Proteomic analysis of HLA-I complexes has been used to identify antigenic peptides from known genes, and these experiments rely on immunoprecipitation of HLA complexes with bound peptides^63^. We reasoned that HLA-I immunoprecipitation serves as an ideal enrichment step to enhance microprotein peptide detection and simultaneously allow for identification of microprotein-derived antigens (Fig. 9a). Searching a published HLA-I proteomics dataset^64^ using the human Swiss-Prot database appended with the 7,554 novel smORF-encoded microproteins, we identified peptides from 320 microproteins (Fig. 9b). Of these, 192 were from smORFs identified in at least two Ribo-Seq experiments (Supplementary Data 3), and 41 lacked an in-frame AUG start codon. A previous study was also able to detect over 100 microprotein peptides in the same proteomics dataset, which is consistent with and expanded by these data^65^. Representative spectra of peptides from smORF-encoded microproteins demonstrate good fragment ion coverage, regardless of the number of times detected and cell lines found in by Ribo-Seq (Fig. 9c). We then validated the binding of three peptides to the HLA-I complex using a fluorescence-based competition assay and observed clear displacement of the HLA-I reference peptide by all three (Fig. 9d and Supplementary Fig. 8). Thus, we validated the translation of hundreds of smORFs at the protein level and obtained evidence that they are capable of being presented on HLA-I complexes like annotated proteins.

**Figure 9.**
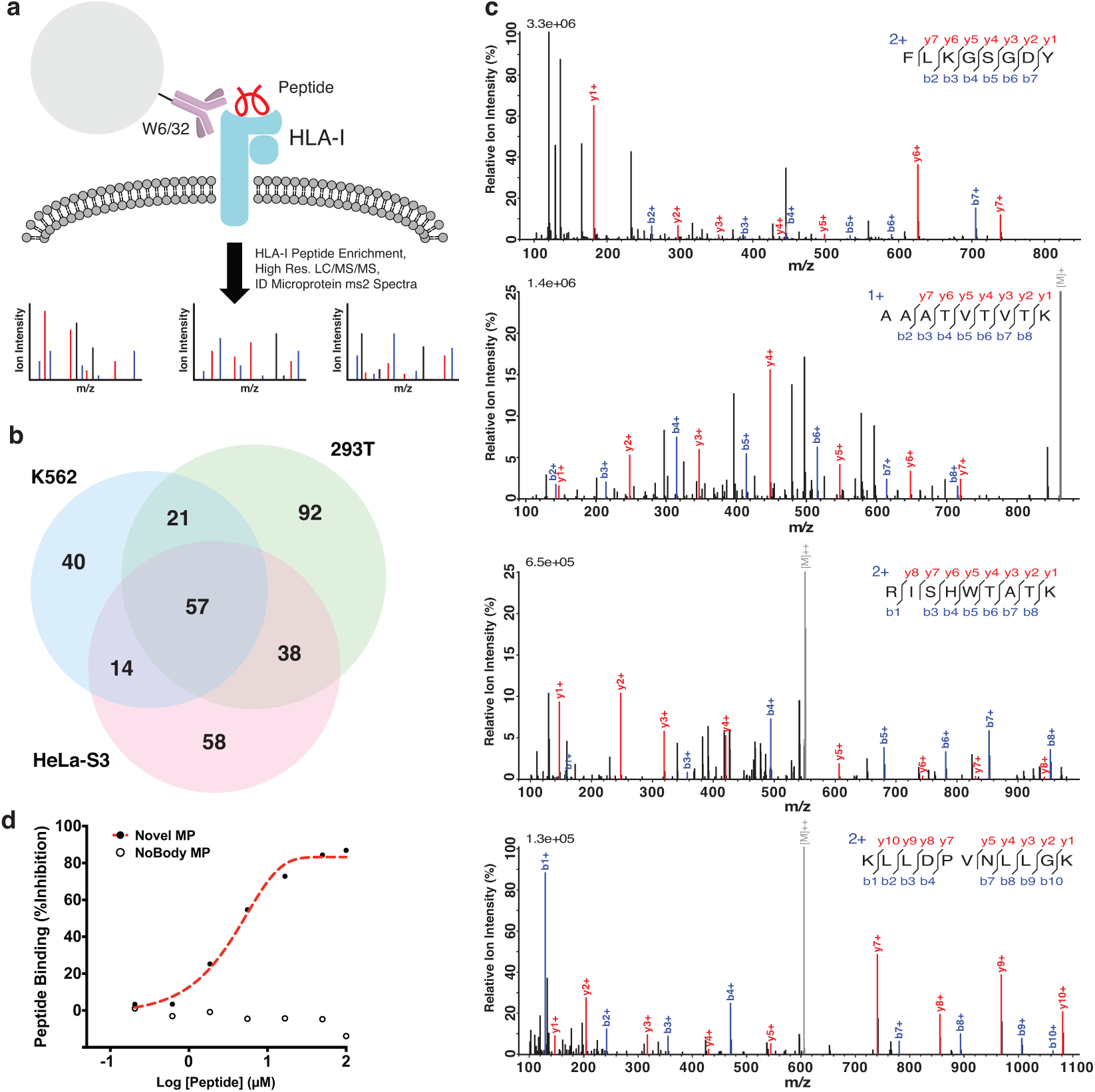
Novel microproteins were detected in HLA-I complexes. **a** Schematic of HLA-I bound peptide enrichment experiment carried out in Bassani-Sternberg et al.^64^. The pan-HLA-I antibody, W6/32, was used to pull-down and enrich HLA-I complexes, and bound small peptides were further enriched by solid phase extraction. High resolution tandem mass spectrometry data of enriched HLA-I peptide samples (PXD000394) was then searched against a database containing human Uniprot proteins and the 7,554 novel smORF-encoded microproteins. **b** 320 novel microproteins were identified across all three cell lines, of which 130 (41%) were identified in at least two Ribo-Seq experiments and therefore high confidence. **c** ms2 spectra examples (top-bottom) for smORFs found in: three cell lines, two cell lines, one cell line (multiple experiments), and one cell line (single experiment). **d** Binding of a novel microprotein peptide, RMKDFLCLK (chr1:39875291-39875422), was validated by a competition-based fluorescence assay^74^. The novel microprotein peptide was able to compete off the control peptide, indicating binding, while the negative control peptide, TPNGGSTTL, from the recently characterized microprotein NoBody^16^ was unable to.

## Discussion

Utilizing our top-down smORF annotation workflow, we were able to rigorously annotate thousands of novel protein-coding smORFs across 16 Ribo-Seq experiments in three human cell lines. By analyzing individual experiments, we showed that predicting smORF translation from Ribo-Seq data is noisier than for annotated genes (Fig. 2b,c and Supplementary Data 2). Differences in Ribo-Seq resolution, sequencing depth, and variability in sequencing library construction as well as biological variations such as passage number and cell density play significant roles in smORF translation analysis. However, given that annotated genes were also subjected to these effects and yet had much greater overlap, it is most likely that overall lower translation levels explain why smORFs are more difficult to detect reproducibly. We also show that it is beneficial to use a range of RNase I digestion conditions to annotate smORFs, as there are several hundred reproducibly detected smORFs that were only identified in lower resolution or higher resolution datasets (Supplementary Data 2). Importantly, we demonstrate that *de novo* transcriptome assembly is necessary for comprehensive smORF annotation (Fig. 7 and 8).

While our data represent a significant step in comprehensive protein-coding smORF annotation, we expect future studies to find additional novel smORFs. First, these numbers are an underestimation because we chose to exclude smORFs that overlap with longer ORFs in our analyses, though such smORFs are known^22^. By definition, overlapping smORFs have RPF reads aligned out-of-frame relative to another ORF which limits the scoring of both, especially for smORFs with a high percentage of overlap or a low abundance relative to the other ORF. Our highest resolution datasets may be suitable for identifying abundant overlappers, however, we expect to find a significant number of artifacts using our lower resolution datasets due to the higher percentage of noisy out-of-frame reads. Second, we utilized ENCODE cell lines, which are valuable but are likely different from primary cells or tissues. Future studies would benefit from including more physiologically relevant samples to determine if the smORFs we detect rely on the cellular context. Finally, improvements to sample preparation methods, such as long read sequencing for transcript assembly and small RNA library construction, and to computational methods for short read alignment and analysis of RPFs for translation will be critical for complete annotation of functional smORFs.

For many smORFs, these data provide the first evidence of translation. Therefore, we propose using reproducibility as one filter for follow-up functional studies. Over 2,500 smORFs were called translated in multiple experiments across all cell lines and thus are higher confidence annotations. More confident still are those smORFs found in multiple cell lines, because in order to do so both the transcript and Ribo-Seq evidence must be reproducible. Being found in multiple cell lines also suggests that these smORFs have a general cellular function, which might explain their increased canonical AUG start codon usage (Fig. 4a). Nevertheless, smORFs identified in a single experiment are worth including in large scale studies, as many of these just failed the stringent RibORF scoring filter in other experiments and might pass with higher sequencing depth or in a differently prepared sample. Supporting this hypothesis, we were able to detect peptides from singly identified smORFs in HLA-I proteomics datasets (Fig. 9c and Supplementary Data 3). Furthermore, there were hundreds of annotated genes that were only detected in a single experiment (Fig. 2b).

Beyond reproducibility, useful methods for uncovering biologically functional smORFs include identifying those that are regulated, bound to protein complexes, or evolutionarily conserved. For example, we found dozens of novel smORFs that were regulated at the transcription level and one regulated at the translation level during ER stress (Fig. 3b,d). We also detected hundreds of microprotein peptides bound to HLA-I complexes in a variety of human cell lines (Fig. 9). Expression regulation and detection by mass spectrometry further validate these smORFs and position them well for functional characterization studies. For the microprotein-derived antigens, the next important step will be to test if any of these are immunogenic. Functional inferences can also be drawn from microprotein sequence conservation (Fig. 4d-f), as several characterized smORFs have excellent conservation by PhyloCSF and BLAST^13–17^. Having identified thousands of smORFs, additional biological data can easily be mined to elucidate their roles.

This study serves two key purposes, the development of a refined workflow for smORF annotation and the curation of a human smORF database for functional characterization. Given the sheer number of protein-coding smORFs annotated, their diversity in amino acid composition, and cell type specificity, we anticipate smORFs being involved in all facets of biology. In addition, new insights into translational regulation can be gained by studying polycistronic RNAs (Fig. 6c,d and Supplementary Fig. 7) and how multiple start sites are employed for the same reading frame (Fig. 5). These results also add to the growing evidence that some ncRNAs might operate as both a functional molecule and a coding template. In summary, smORFs offer a rich opportunity for uncovering new biology, and in the future perhaps a new avenue for therapeutic discovery.

## Methods & Materials

### Cell Culture

HeLa-S3 cells were purchased from ATCC (CCL-2.2). HEK293T cells were purchased from GE Life Sciences (HCL4517). K562 cells were purchased from Sigma-Aldrich (89121407). HEK293T, and HeLaS3 cells were maintained in DMEM (Corning, 10-013-CV) supplemented with 10% Fetal Bovine Serum (FBS; Corning, 35-010-CV). K562 cells were maintained in RPMI 1640 (Corning, 10-040-CV) supplemented with 10% FBS. All cells were maintained at 37 ºC with 5% CO_2_.

### Paired-End RNA-Seq and *de novo* Transcriptome Assembly

The HEK293T Cufflinks assembled transcriptome was generated previously^66^, and used to create the ORF database for scoring translation with RibORF. For HeLaS3 and K562, total RNA was harvested and purified from two biological replicates using an RNeasy Kit (Qiagen) with gDNA eliminator columns. For each cell line, two separate cDNA libraries were prepared for each replicate: one using the TruSeq Stranded mRNA Kit (Illumina) and the other using the TruSeq Total RNA Kit (Illumina). This allowed for representation from poly-A tailed mRNA and non-poly-A RNAs in the transcriptome assembly. Paired-end 125 or 150 base reads were collected for all 4 libraries on a single lane of an Illumina HiSeq 2500 or NextSeq 500, respectively. At least 250M reads were generated for each cell line. Aligned reads were assembled into transcripts by Cufflinks using default parameters, fragment bias correction, multi-read correction, fr-firststrand library construction, and the hg19 human genome sequence as a guide.

### Ribosome Footprinting

Preparation of ribosome footprints for Ribo-Seq experiments was performed as described^30^ with some modifications. For all ribosome footprinting experiments, adherent cells were grown to about 80% confluency in 10 cm or 15 cm diameter tissue culture dishes and suspension cells were grown to a density of approximately 500,000 cells/mL. Cells were washed with 5 mL ice-cold Phosphate Buffered Saline (PBS) with 100 µg/mL cycloheximide (CHX) added. Immediately after removing PBS, 400 µL of ice-cold lysis buffer (20 mM Tris-HCl, pH 7.4, 150 mM NaCl, 5 mM MgCl_2_, 1% Triton X-100, with 1 mM DTT, 25 U/mL Turbo DNase (Thermo Fisher, AM2238), and 100 µg/mL CHX added fresh) was dripped onto the plate or added to the cell pellet. Cells were incubated on ice in lysis buffer for 10 min with periodic vortexing and pipetting to disperse the cells. The lysate was then clarified by centrifugation at 15,000 *g* for 10 min. Cell lysates were flash frozen and liquid nitrogen and stored at −80ºC for up to 5 d prior to ribosome footprinting. For experiments profiling translation initiation, the same procedure was followed except for the addition of either 2 µg/mL harringtonine (abcam) for 2 min or 20 µg/mL lactimidomycin (Calbiochem) for 30 min to media prior to PBS wash and lysis. A variety of digestion conditions were tested in this study and are summarized in the Supplemental Methods. Briefly, RNA digestions using 250 U RNase I (Thermo Fisher, AM2294) per 100 µL lysate were used in the low resolution 293T and HeLaS3 experiments. For high-resolution experiments, 15 to 30 U TruSeq Nuclease (Illumina) was used to digest 30 to 60 µg RNA in up to 300 µL lysate. Digestion reactions were run for 45 to 60 min at RT and quenched with 100 to 200 U Superase-In RNase I inhibitor (Thermo Fisher) on ice. Following digestion, ribosome protected fragments (RPF) were purified from small RNA fragments using MicroSpin S-400 HR columns (GE Life Sciences) according to the TruSeq Ribo Profile Kit (Illumina). Low resolution experiments were cleaned up with Zymo RNA Clean & Concentrator-25 kit, while high resolution experiments were purified by acid phenol:chloroform extraction followed by isopropanol precipitation. Ribosomal RNAs were depleted from RPF fragments by Ribo-Zero Mammalian Kit (Illumina) following the manufacturer’s protocol. cDNA sequencing libraries were then prepared using the TruSeq Ribo Profile Kit (Illumina) following the manufacturer’s protocol. Single-end 50 base reads were collected for each library on an Illumina HiSeq2500 with no more than 4 samples sequenced on a single lane. Each Ribo-Seq experiment was prepared from a different biological replicate except for K562 HiRes1 & 2 which were prepared from the same lysate using different digestion conditions. For K562 HiRes3, CHX was added to the media prior to pelleting cells and washing with PBS.

### Ribo-Seq and Short Read RNA-Seq Read Processing

Ribo-Seq and accompanying short fragment total RNA-Seq reads were first trimmed of excess 3’ adaptor sequences as in Calviello et al.^30^ using the FASTX-toolkit. Trimmed Ribo-Seq reads aligning to tRNA and rRNA sequences were then removed using STAR v2.5.2b^67^ as in Wang et al.^68^. Next, the remaining Ribo-Seq reads were aligned to the UCSC hg19 human genome assembly containing chromosomes 1-22, X, and Y with the hg19 refGene transcript annotation using STAR. Up to two mismatches were allowed during alignment, keeping only uniquely mapped reads. Ribo-Seq and RNA-Seq alignments were checked for overall quality using the CollectRnaSeqMetrics script from the Picard Tools software suite.

### RibORF Scoring

Following Ribo-Seq read processing and quality control, the RibORF software package^38^ was used to score individual ORFs for translation. First, metagene analysis was conducted using coding genes from the hg19 refGene annotation included with RibORF. Metagene analysis is run for individual processed read lengths ranging from 25-34 nt. Using the metagene plots, the offset shift needed to align the 5’-most position with the A-site, or +3 position, for each read length is assessed. Next, the entire Ribo-Seq alignment is corrected by the offset shift for each length. For high-resolution data, reads ranging from 25 to 30 nt in length were included depending on the sample’s footprint length distribution. For lower resolution data, reads ranging from 28 to 35 nt were included. The offset-corrected read alignments were used for scoring individual ORFs as translated. Following the suggestions of the RibORF developers, only ORFs with RibORF scores ≥0.7 and at least 10 reads mapped to the ORF were considered translated in each individual Ribo-Seq dataset. Each Ribo-Seq dataset was analyzed individually for translated smORF predictions. RNA coverage and Ribo-Seq A-site plots for individual smORFs were plotted using R scripts.

### Defining ORFs

RibORF does not define boundaries of putative ORFs based on Ribo-Seq coverage and thus requires a user-generated list of candidate ORFs. Generation of ORF databases from the *de novo* assembled transcriptome of each cell line was done using a custom java script, GTFtoFASTA (Supplementary Data 4). For each cell line’s *de novo* assembled transcriptome, ORFs were defined by identifying the most distal in-frame upstream AUG start codon for every stop codon across all three reading frames. Because Ribo-Seq evidence is expected to occur solely within a putative ORF, it is important to limit ORFs to AUG start codons, which are mostly likely to be initiation sites based on the scanning model of translation, when available instead of beginning at upstream stops. However, if no AUG start codon is found, the ORF was defined from stop codon to stop codon to allow for the identification of non-AUG initiated smORFs. The resulting millions of ORFs were then assembled into a database containing the exon coordinates for each ORF in refFlat format. In Ribo-Seq datasets, translation termination peaks are often overrepresented and have a higher fraction of reads aligned to the second position (out of frame) compared to non-stop codons, as observed by metagene analysis (Fig. 2a). Therefore, for RibORF scoring, only the first position of the stop codon was included in the ORF as opposed to the full stop codon. By only including the first position of the stop codon in the ORF definition, we limited the scoring penalty that frequently occurs due to the higher frequency of out of frame reads. A previous study dealt with the extreme nature of translation termination peaks by excluding the stop codon altogether from scoring^27^, while others include the entire stop codon and do not handle it differently^38^. While the majority of smORFs called translated do not change whether the stop codon is included or not, our strategy results in the highest number of predicted protein-coding smORFs and offers the best overlap with each alternative option across all different levels of overall Ribo-Seq resolution tested (Supplementary Fig. 8).

### Differential Translation Analysis

Differential translation analysis was conducted using the R package Xtail v1.1.5^50^. First, HTSeq-count^69^ in intersection-strict mode was used to calculate total RNA read counts for hg19 refGene annotations. For smORFs, HTSeq-count was run in union mode and allowed for non-unique reads to be counted. RPF read counts for the same annotations were calculated using the custom python script in Xiao et al.^50^, which retains only uniquely mapped reads occurring within the middle of the CDS region. For hg19 RefGene annotated genes, reads aligning after the first 15 codons and before the last 5 codons were counted. For novel protein-coding smORFs, reads aligning after the first and before the last codon were counted. Xtail was used to calculate the log2 fold-changes in translation efficiency (TE) between DMSO- and tunicamycin-or thapsigargin-treated cells from the read count tables. Genes not considered ‘stable’ by xtail and with a log2 fold-change ≥ 1 or ≤ −1 were assigned as either ‘homodirectional,’ ‘transcription-only,’ or ‘translation-only’ category of differential translation. DESeq2^70^ was also run in parallel with Xtail to calculate differential mRNA expression for hg19 refGene annotations and smORFs. Plots summarizing the results from both analyses were generated using R.

### PhyloCSF and BLAST Analyses of protein-coding smORFs

Smoothed PhyloCSF scores for the 29-mammals alignment were extracted for all smORFs from the UCSC genome browser’s PhyloCSF Track Hub using the bedtools map function. The scores represent the log-odds that codons in the smORF are in the coding state. The average smoothed PhyloCSF scores are shown for each protein-coding smORF by exon (Supplementary Data 2).

All smORFs were queried for similarity against the non-redundant database using tBLASTn and BLASTP under default parameters. BLAST alignments were considered significant if the BLAST score ≥ 80 or if ≥80% of the microprotein sequence matched ≥80% of the aligned subject sequence. This second condition allowed for the identification of short but high similarity sequence alignments, which otherwise have a low BLAST score under default parameters.

### Mass Spectrometry Data Analysis

Mass spectrometry data from PXD000394^64^ were downloaded from the PRIDE archive. Tandem mass spectra were extracted from RAW files using RawConverter 1.0.0.0. Next, the spectra were searched against a database containing human Swiss-Prot proteins, novel microproteins, and common contaminants using ProLuCID^71^. The enzyme specificity was set to none and no variable modifications were included. The false discovery rate was set to 1% for peptides. Identified spectra were then filtered and grouped into proteins using DTASelect^72^. Mass spectrometry analyses were separated by different cell lines from the study. We also utilized the pFind 3 Open-pFind^73^ search engine to identify microprotein-derived peptides by an open search strategy, which allows for many variable modifications, using the same database and false discovery rate.

### HLA-I peptide binding assay

The affinities of novel microprotein-derived peptides for HLA-I were measured as previously described^74^. Briefly, SupB15 cells (HLA-I: A3, A11, B51, B52 serotype) were harvested and the cell surface HLA complex was disassembled by treating with citric acid elution buffer (pH 2.9) for 90 seconds. Then, cells were incubated with a high-affinity fluorescein-labeled reference peptide KVFPC(FITC)ALINK (1 μM) and increasing concentrations of a non-labeled microprotein-derived peptide for 20 hours at 4°C. A negative control peptide from the recently characterized microprotein NoBody^16^ (TPNGGSTTL, B7 serotype binder) was also tested for comparison. Fluorescence intensities were measured by flow cytometry. Binding of novel microprotein-derived peptides at each concentration was calculated as percentage inhibition of reference peptide binding relative to background (without reference peptide, MF_bg_) and the maximal response (reference peptide only, MF_ref_) using the following equation: Inhibition (%) = (1 – (MF – MF_bg_)/(MF_ref_ – MF_bg_))*100

The data were then plotted and fit for IC50 calculation using Prism 5.

### Peptide synthesis

Peptides were purchased from Peptide 2.0. Fluorescein-labeled reference peptide KVFPC(FITC)ALINK was synthesized by covalently coupling of fluorescein to the cysteine residue with 5-(iodoacetamido)fluorescein (Marker Gene Technologies, M0638) for use in the HLA-binding assay. All peptides were purified by high-performance liquid chromatography and confirmed by mass spectrometry.

## Data Availability

All sequencing datasets generated in this study are available through GEO (GSE125218).

## Acknowledgements

We thank the Saghatelian lab for helpful comments and suggestions throughout the study, and Prof. Nicholas Ingolia for advice on RNase I digestion conditions. We also thank Manching Ku, Nasun Hah, and the Salk Institute Next Generation Sequencing Core for preparation of RNA-Seq libraries and high-throughput sequencing of Ribo-Seq and RNA-Seq libraries. This research was supported by NIH/NIGMS (R01 GM102491, A.S.), Leona M. and Harry B. Helmsley Charitable Trust grant (A.S.), Dr. Frederick Paulsen Chair/Ferring Pharmaceuticals (A.S.), NIH/NIGMS postdoctoral fellowship (F32 GM123685, T.F.M.), George E. Hewitt Foundation for medical research (Q.C.), Pioneer Fellowship (D.T.). This work was also supported by the Razavi Newman Integrative Genomics and Bioinformatics Core and the Next Generation Sequencing Core Facilities of the Salk Institute with funding from the NIH-NCICCSG (P30 014195) and the Chapman Foundation.

## Author contributions

T.F.M. and A.S. conceived the project, designed the experiments, and wrote the manuscript. T.F.M. performed cell culture, and prepared RPFs and total RNA. T.F.M. and C.D. prepared Ribo-Seq libraries. T.F.M. analyzed Ribo-Seq and RNA-Seq data, developed the smORF annotation workflow, and wrote custom scripts to generate Ribo-Seq plots. M.N.S. performed *de novo* transcriptome assembly and generated ORF databases. Q.C. performed HLA-I experiments. T.F.M. and D.T. analyzed HLA-I proteomics data. All authors discussed the results and edited the manuscript. A.S. supervised the study.

---

**Supplementary Figure 1.**
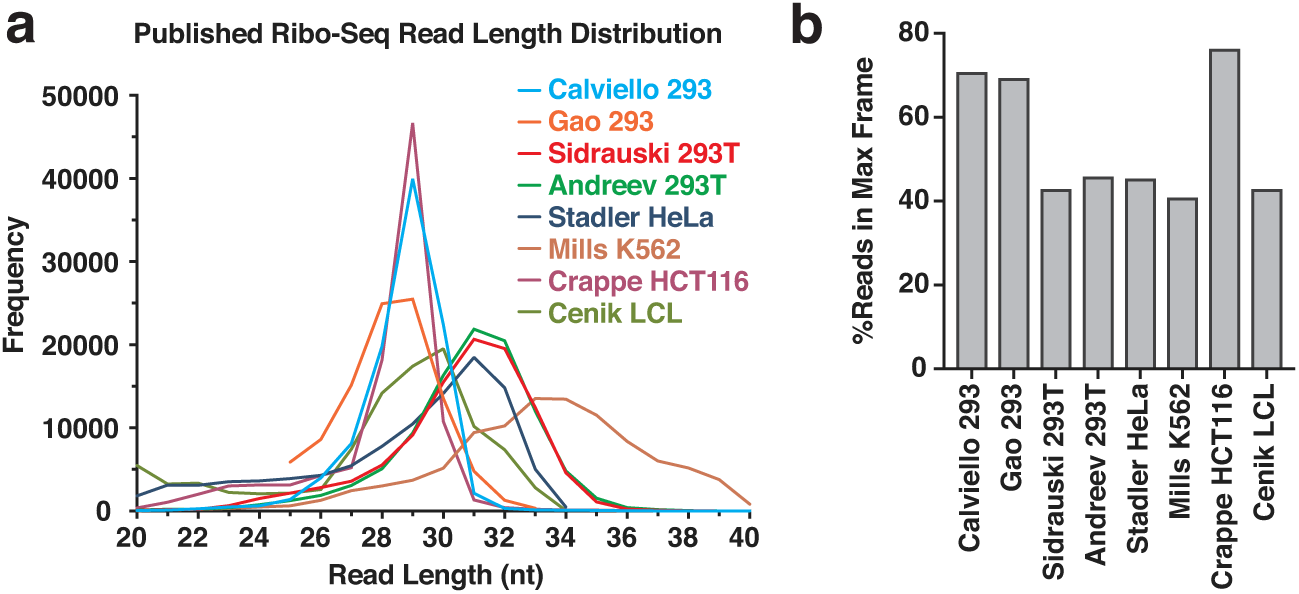
Published Ribo-Seq datasets show a wide range of overall resolutions. **a** Ribo-Seq datasets from 8 published studies^30,40,41,55,62,76-78^ using 6 different cell lines were processed and mapped to hg19 using the same pipeline as used for our datasets. 100,000 random reads were sampled to determine the frequency distribution of footprint lengths ranging from 20-40 nt. Ribosome footprint read length distributions from these datasets vary widely. Three datasets have distributions which peak in the ideal 28-29 nt footprint size, indicating complete digestion of unprotected RNA, while the other distributions are broader and peak in 31-34 nt range. **b** Metagene analysis was performed and the average percentage of reads aligned to the coding reading frame was calculated as a measure of the overall 3-nt periodicity and resolution of the datasets. Datasets which had ribosome footprint lengths peaking at the ideal 28-29 nt size had >70% of reads in-frame with the coding region, which is high overall resolution, while the other datasets had <50% reads in-frame.

**Supplementary Figure 2.**
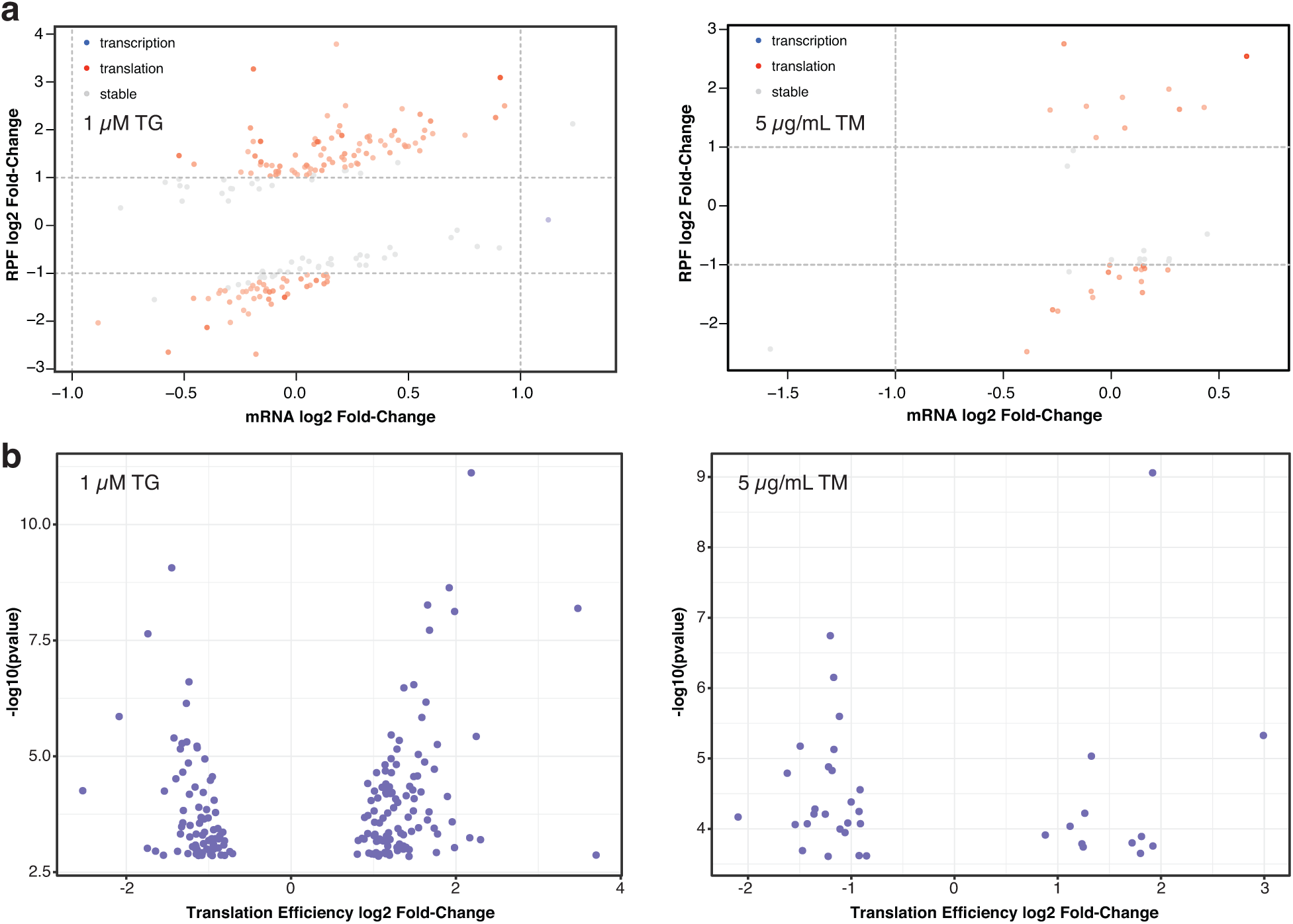
Dozens of annotated genes are translationally regulated in response to ER stress. **a** Scatter plots showing the log2 fold changes in normalized RPF (Ribo-Seq, y-axis) and mRNA (RNA-Seq, x-axis) read counts for RefSeq annotated genes in cells treated with either 1 µM thapsigargin (TG, left) or 5 µg/mL tunicamycin (TM, right) relative to DMSO-treated cells. Only genes with significant changes (p_adj_ < 0.1) in translation efficiency (TE) are plotted. Genes with TE log2 fold changes ≥ 1 or ≤ 1 and RPF or mRNA log2 fold changes ≥ 1 or ≤ 1 are colored. Large changes in TE are colored red if driven predominantly by changes in translation and blue if driven by changes in transcription. Genes colored grey either do not have large enough changes in TE or lack large changes in both translation and transcription and are therefore considered relatively stable. **b** Volcano plots showing the −log10(p value) and TE log2 fold change for RefSeq annotated genes in cells treated with either TG (left) or TM (right) relative DMSO-treated cells. Only genes with significant changes (p_adj_ < 0.1) in translation efficiency (TE) are plotted.

**Supplementary Figure 3.**
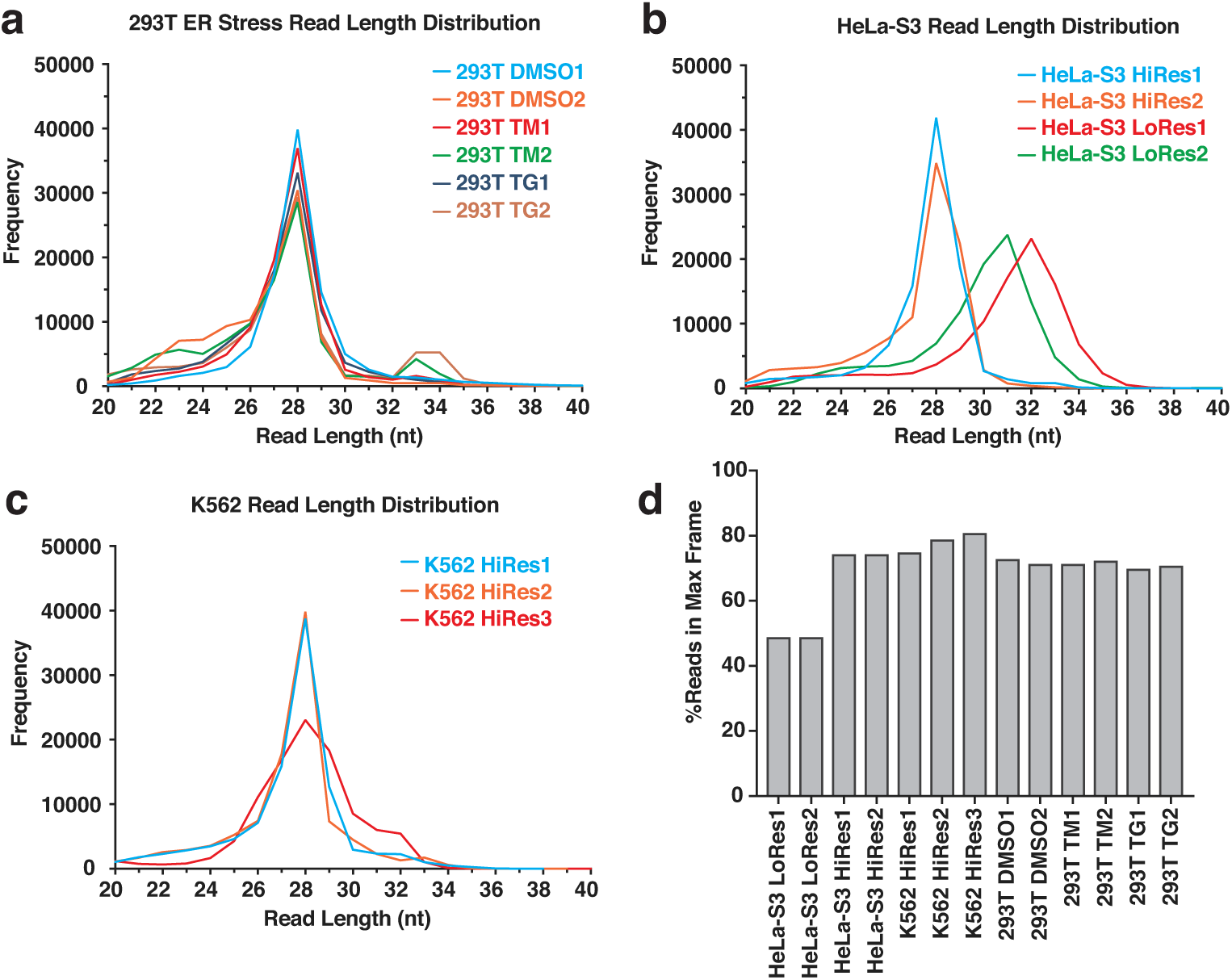
Ribosome footprint distributions and 3-nt periodicity measurements for HeLa-S3, K562, and drug-treated HEK293T samples. **a** Ribosome footprint read length distributions for HeLa-S3 datasets. Two different digestion protocols were employed resulting in 2 datasets peaking at 28 nt with a narrow distribution and 2 datasets peaking at 31-32 nt with a broader distribution. **b** Ribosome footprint read length distributions for K562 datasets. Three different digestion protocols were employed with all 3 datasets peaking at 28 nt. HiRes3 shows a broader distribution than HisRes1 and HiRes2. **c** Ribosome footprint read length distributions for drug-treated HEK293T datasets. All samples were prepared using the same digestion protocol resulting in narrow 28 nt peaks. **d** Metagene analysis was performed and the average percentage of reads aligned to the coding reading frame was calculated. Datasets which had ribosome footprint lengths peaking at the ideal 28-29 nt size had >70% of reads in-frame with the coding region, which is high overall resolution, while the other datasets had ~50% reads in-frame.

**Supplementary Figure 4.**
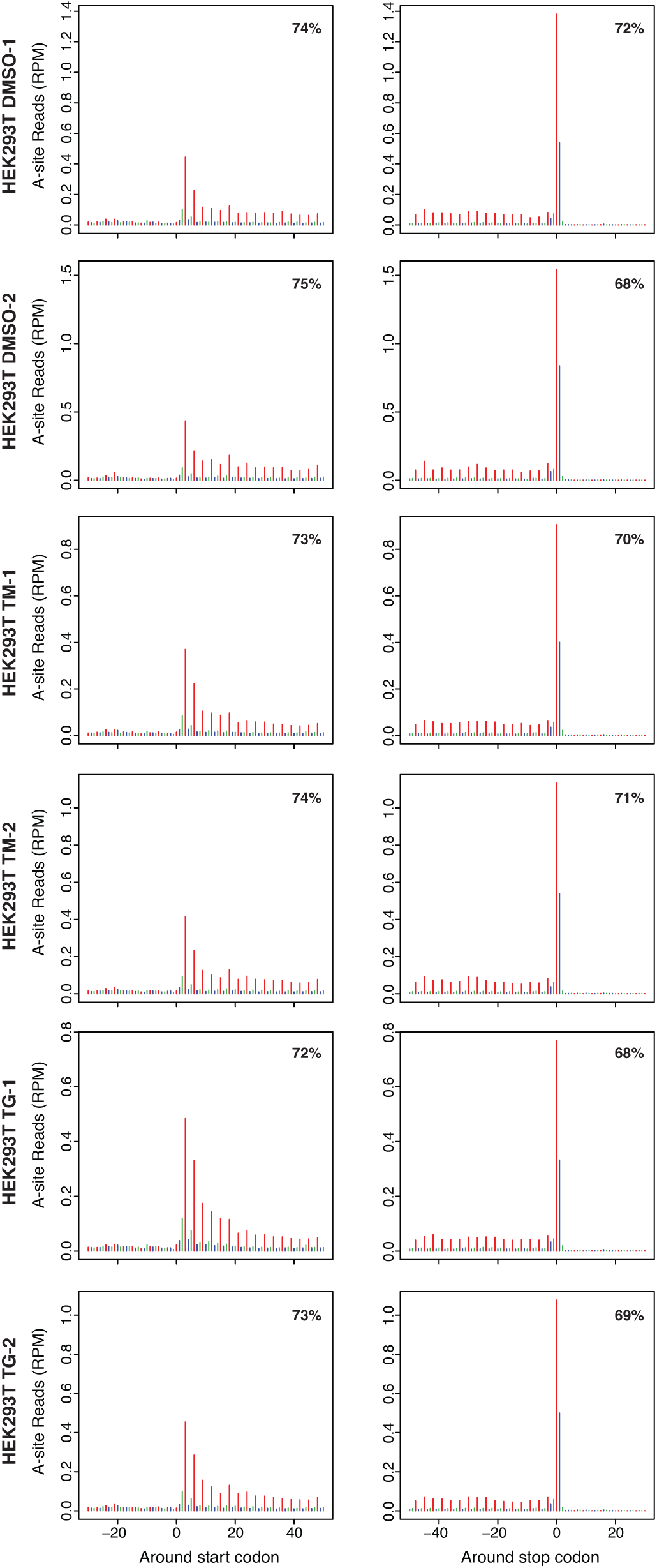
Metagene plots for drug-treated HEK293T Ribo-Seq datasets. Metagene plots showing RPF read alignment around the start site and stop site for each treated HEK293T replicate. 25-29 nt reads were used for all datasets.

**Supplementary Figure 5.**
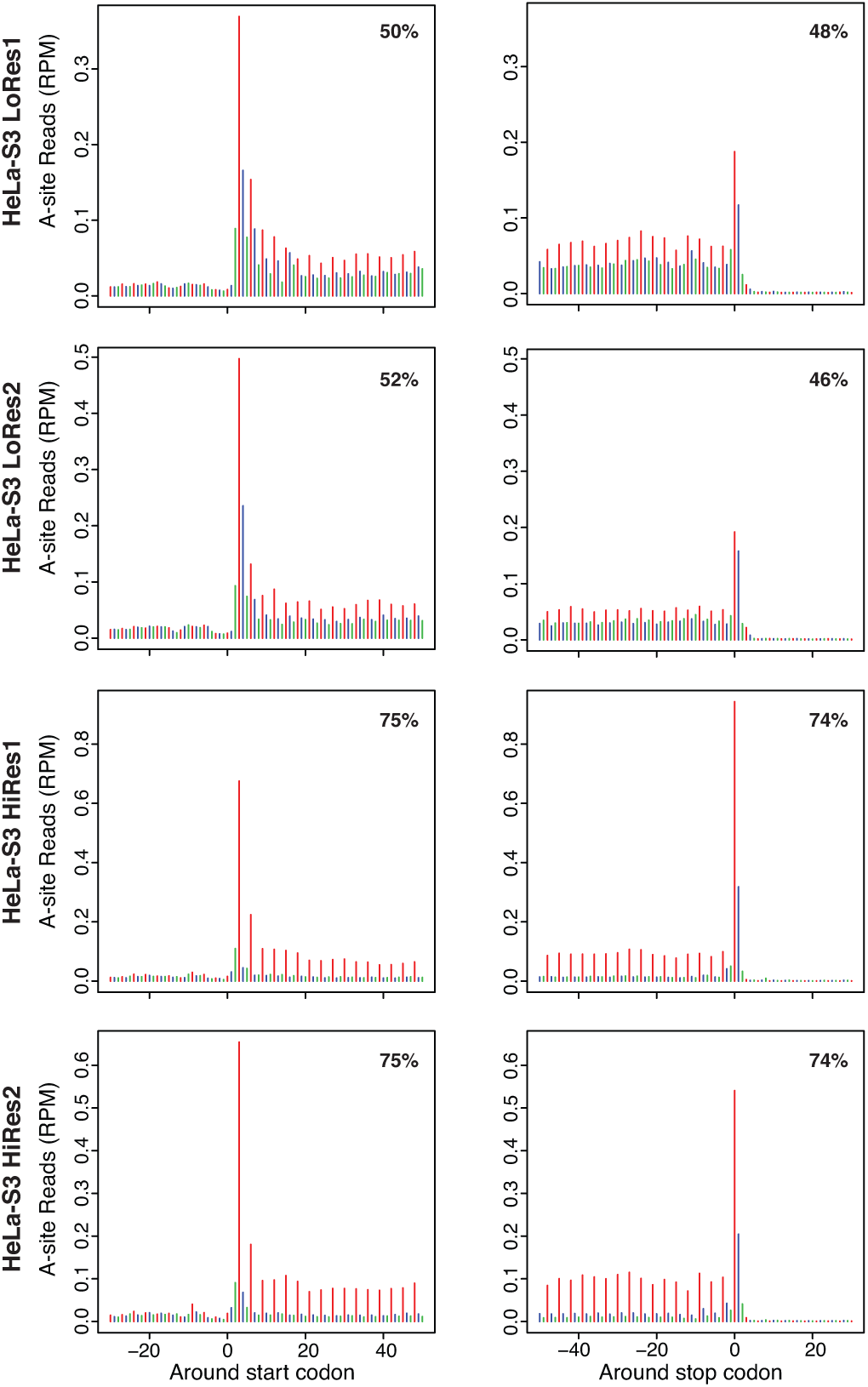
Metagene plots for HeLa-S3 Ribo-Seq datasets. Metagene plots showing RPF read alignment around the start site and stop site for each HeLa-S3 replicate created using RibORF. The 5’-position of each RPF read was shifted to the ribosomal A-site and then mapped to all hg19 RefSeq coding transcripts, which were used to construct the metagene. The metagene coding region is aligned to frame 1 (red), and frame 2 (blue) and frame 3 (green) are out of frame. The percentage of reads aligned to the coding region is noted in the top corner. 31-35 nt reads were used for LoRes1, 29-33 nt for LoRes2, and 25-29 nt for both HiRes1 and HiRes2.

**Supplementary Figure 6.**
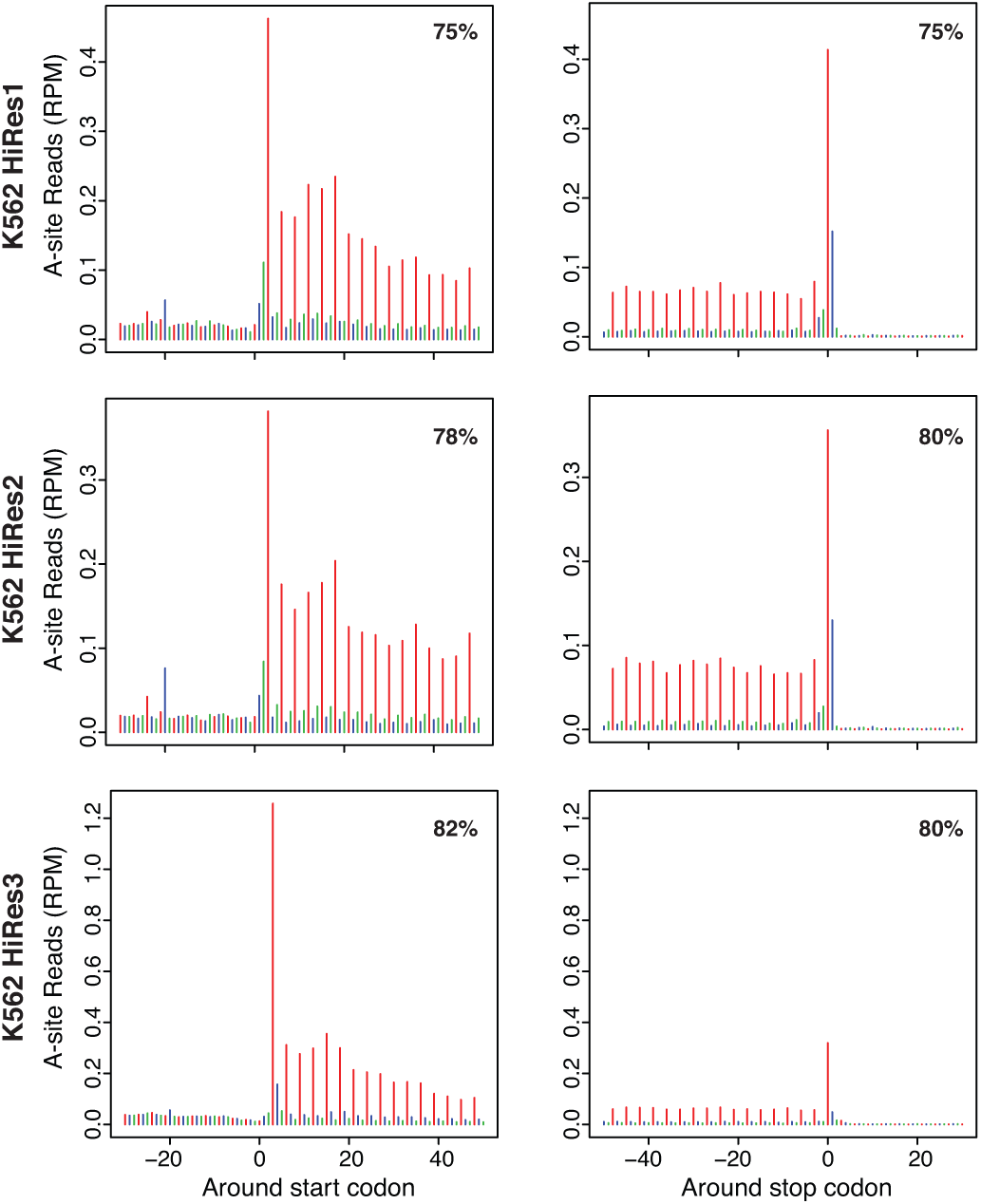
Metagene plots for K562 Ribo-Seq datasets. Metagene plots showing RPF read alignment around the start site and stop site for each K562 replicate. 25-29 nt reads were used for both HiRes1 and HiRes2, and 25-30 nt for HiRes3.

**Supplementary Figure 7.**
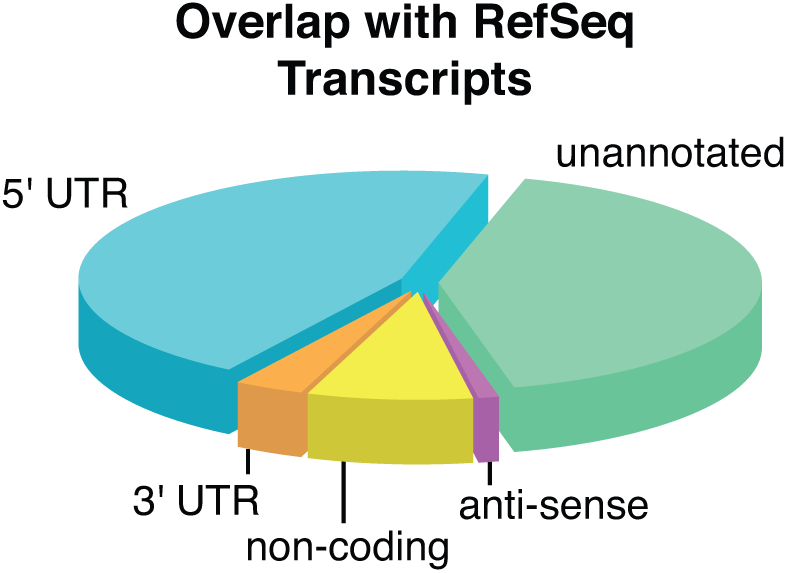
smORF locations in transcriptome. Pie chart showing the locations of all novel smORFs relative to annotated RefSeq transcripts.

**Supplementary Figure 8.**
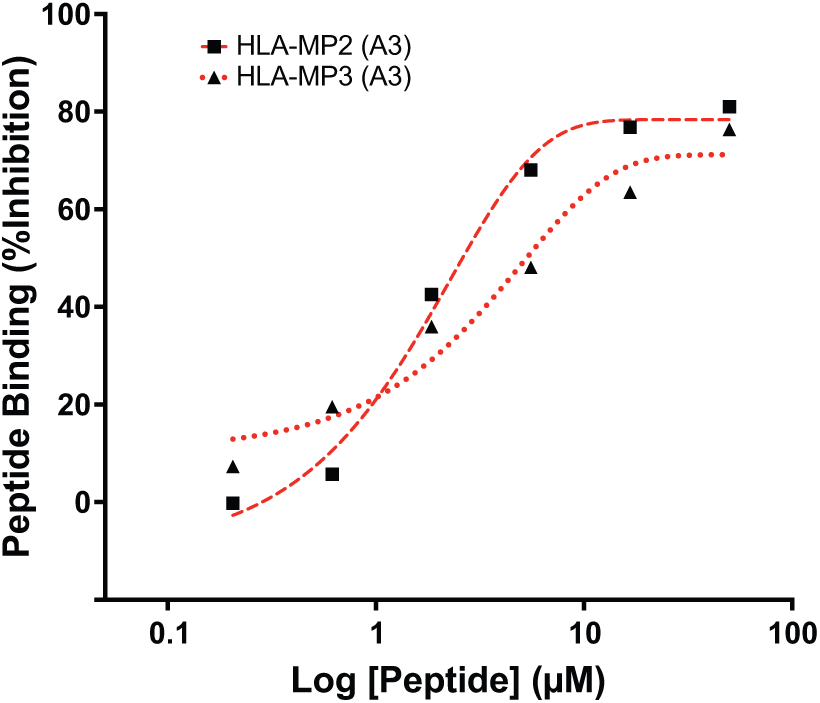
Validation of novel microprotein peptides binding to HLA. **I.** Binding of two additional novel microprotein peptides identified in a published HLA-I proteomics dataset^64^, MTMSTILSKK (chr5:8460071-8460175, HLA-MP2) and HMMDKRLGEK (chr21:46710407-46710529, HLA-MP3), was validated by a competition-based fluorescence assay^74^. Both novel microprotein peptides were able to compete off the control peptide, indicating binding.

**Supplementary Figure 9.**
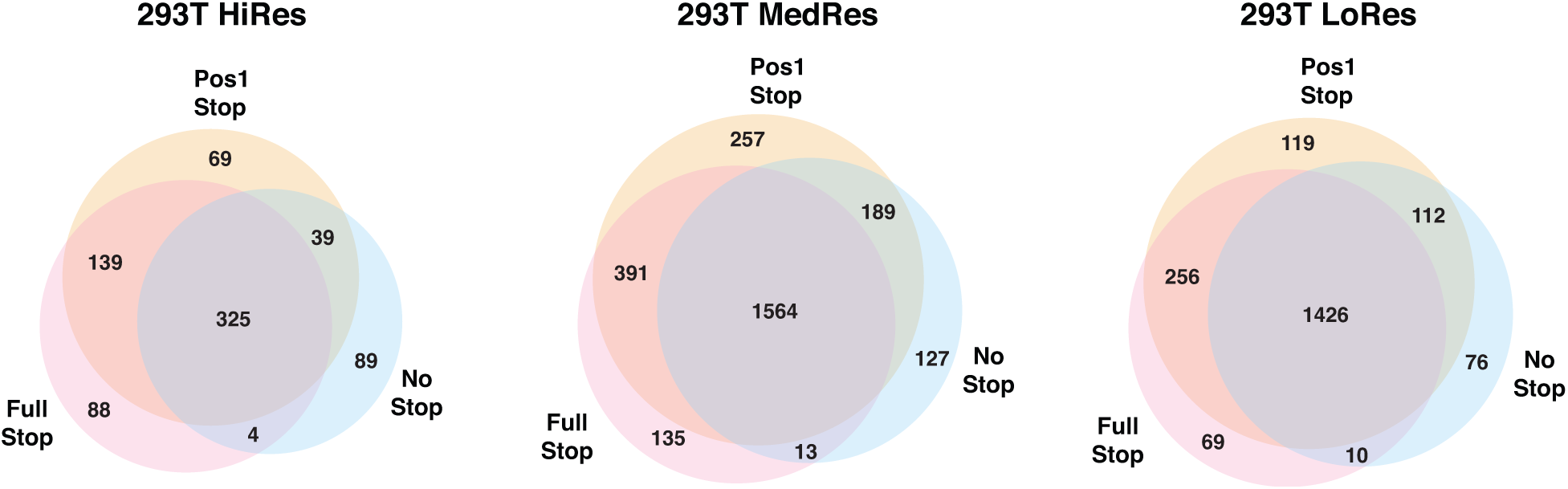
Effect of stop site inclusion on smORF prediction. When analyzing Ribo-Seq reads shifted to the ribosomal A-site, the final codon with ribosome coverage is the stop site. This codon is often unique in that it usually has enriched read coverage compared to codons in the middle of the ORF. In addition, the second position of the stop codon is often enriched relative to middle codons, which can affect scoring methods utilizing the percentage of reads in-frame. The effect of including the entire stop site (Full Stop), just the first position of the stop site (Pos1 Stop), or no stop site at all (No Stop) on the number smORFs called translated using our pipeline was tested across our HEK293T datasets of varying resolution and sequencing depth. In all datasets tested, most smORFs are called translated regardless of whether all or none of the stop codon is included. However, the overall number of smORFs called translated was highest in every case when including only the first position of the stop site. Using only the first position also offered the best balance for the MedRes and LoRes datasets as more hits from “full stop” and “no stop” were captured than lost.

